# Slow oscillation-spindle coupling strength predicts real-life gross-motor learning in adolescents and adults

**DOI:** 10.1101/2021.01.21.427606

**Authors:** Michael A. Hahn, Kathrin Bothe, Dominik P. J. Heib, Manuel Schabus, Randolph F. Helfrich, Kerstin Hoedlmoser

## Abstract

Previously, we demonstrated that precise temporal coordination between slow oscillations (SO) and sleep spindles indexes declarative memory network development (Hahn et al., 2020). However, it is unclear whether these findings in the declarative memory domain also apply in the motor memory domain. Here, we compared adolescents and adults learning juggling, a real-life gross-motor task. We found that improved task proficiency after sleep lead to an attenuation of the learning curve, suggesting a dynamic juggling learning process. We employed individualized cross-frequency coupling analyses to reduce inter and intra-group variability of oscillatory features. Advancing our previous findings, we identified a more precise SO-spindle coupling in adults compared to adolescents. Importantly, coupling precision over motor areas predicted overnight changes in task proficiency and learning curve, indicating that SO-spindle coupling is sensitive to the dynamic motor learning process. Our results provide first evidence that regionally specific precisely coupled sleep oscillations support gross-motor learning.

## INTRODUCTION

Sleep actively supports learning (Diekelmann & Born, 2010). The influential active system consolidation theory suggests that long-term consolidation of memories during sleep is driven by a precise temporal interplay between sleep spindles and slow oscillations (Diekelmann & Born, 2010; Klinzing et al., 2019). Memories acquired during wakefulness are reactivated in the hippocampus during sharp-wave ripple events in sleep (Wilson & McNaughton, 1994; Zhang et al., 2018). These events are nested within thalamo-cortical sleep spindles that mediate synaptic plasticity (Niethard et al., 2018; Rosanova & Ulrich, 2005). Sleep spindles in turn are thought to be facilitated by the depolarizing phase of cortical slow oscillations (SO) thereby forming slow oscillation-spindle complexes during which the subcortical-cortical network communication is optimal for information transfer (Chauvette et al., 2012; Clemens et al., 2011; Helfrich et al., 2019; Helfrich et al., 2018; Latchoumane et al., 2017; Molle et al., 2011; Ngo et al., 2020; Niethard et al., 2018; Schreiner et al., 2021; Staresina et al., 2015).

Several lines of research recently demonstrated that precisely timed SO-spindle interaction mediates successful memory consolidation across the lifespan (Hahn et al., 2020; Helfrich et al., 2018; Mikutta et al., 2019; Molle et al., 2011; Muehlroth et al., 2019). In our recent longitudinal work, we found that SO-spindle coordination was not only becoming more consistent from childhood to late adolescence but also directly predicted enhancements in declarative memory formation across those formative years (Hahn et al., 2020). However, because the active system consolidation theory assumes a crucial role of hippocampal memory replay for sleep-dependent memory consolidation, most studies, including our own, focused on the effect of SO-spindle coupling on hippocampus-dependent declarative memory consolidation. Therefore, the role of SO-spindle coordination for motor learning or consolidation of procedural information remains poorly understood.

While sleep’s beneficial role for motor memory formation has been extensively investigated and frequently related to individual oscillatory activity of sleep spindles and SO (Barakat et al., 2011; Boutin et al., 2018; Fogel et al., 2017; Huber et al., 2004; King et al., 2017; Nishida & Walker, 2007; Pinsard et al., 2019; Tamaki et al., 2013; Tamaki et al., 2008; Vahdat et al., 2017; Walker et al., 2002), there is little empirical evidence for the involvement of the timed interplay between spindles and SO. In rodents, the neuronal firing pattern in the motor cortex was more coherent during spindles with close temporal proximity to SOs after engaging in a grasping motor task (Silversmith et al., 2020). In humans, stronger SO-spindle coupling related to higher accuracy during mirror tracing, a motor adaption task where subjects trace the line of a shape while looking through a mirror (Mikutta et al., 2019). So far, research focused on laboratory suitable fine-motor sequence learning or motor adaption tasks, which has hampered our understanding of memory consolidation for more ecologically valid gross-motor abilities that are crucial for our everyday life (for a review see King et al. (2017)).

Only few studies have investigated the effect of sleep on complex real-life motor tasks. Overnight performance benefits for riding an inverse steering bike have been shown to be related to spindle activity in adolescents and adults (Bothe et al., 2019; Bothe et al., 2020). Similarly, juggling performance was supported by sleep and juggling training induced power increments in the spindle and SO frequency range during a nap (Morita et al., 2012, 2016). Remarkably, juggling has been found to induce lasting structural changes in the hippocampus and mid-temporal areas outside of the motor network (Boyke et al., 2008; Draganski et al., 2004), making it a promising expedient to probe the active system consolidation framework for gross-motor memory. Importantly, this complex gross-motor skill demands accurately executed movements that are coordinated by integrating visual, sensory and motor information. Yet, it remains unclear whether learning of these precisely coordinated movements demand an equally precise temporal interplay within memory networks during sleep.

Previously, we demonstrated that SO and spindles become more tightly coupled across brain maturation which predicts declarative memory formation enhancements (Hahn et al., 2020). Here we expand on our initial findings by investigating early adolescents and young adults learning how to juggle as real-life complex gross-motor task. We first sought to complete the picture of SO-spindle coupling strength development across brain maturation by comparing age ranges that were not present in our initial longitudinal data set. Second, we explicitly tested the assumption that precisely coordinated SO-spindle interaction supports learning of coordinated gross-motor skills.

By leveraging an individualized cross-frequency coupling approach, we demonstrate that adults have a more precise interplay of SO and spindles than early adolescents. Importantly, the consistency of the SO-spindle coupling dynamic tracked the dynamic learning process of a gross-motor task.

## RESULTS

Healthy adolescents (n = 28, age: 13.11 ± 0.79 years, mean ± SD) and young adults (n = 41, age: 22.24 ± 2.15) performed a complex gross-motor learning task (juggling) before and after a full night retention interval as well as before and after a retention interval during wakefulness (**Figure 1**). To assess the impact of sleep on juggling performance, we divided the participants into a *sleep-first* group (i.e. sleep retention interval followed by a wake retention interval) and a *wake-first* group (i.e. wake retention interval followed by a sleep retention interval). Polysomnography (PSG) was recorded during an adaptation night and during the respective sleep retention interval (i.e. learning night) except for the adult *wake-first* group (for sleep architecture descriptive parameters of the adaptation night and learning night as well as for adolescents and adults see **Supplementary file – table 1 & 2**). Participants without prior juggling experience trained to juggle for one hour. We measured the amount of successful three ball cascades (i.e. three consecutive catches) during performance tests in multiple three-minute (min) blocks (3x3 min for adolescents; 5x3 min for adults) before and after the respective retention intervals. Adolescents performed fewer blocks than adults to alleviate exhaustion from the extensive juggling training.

**Figure 1.**
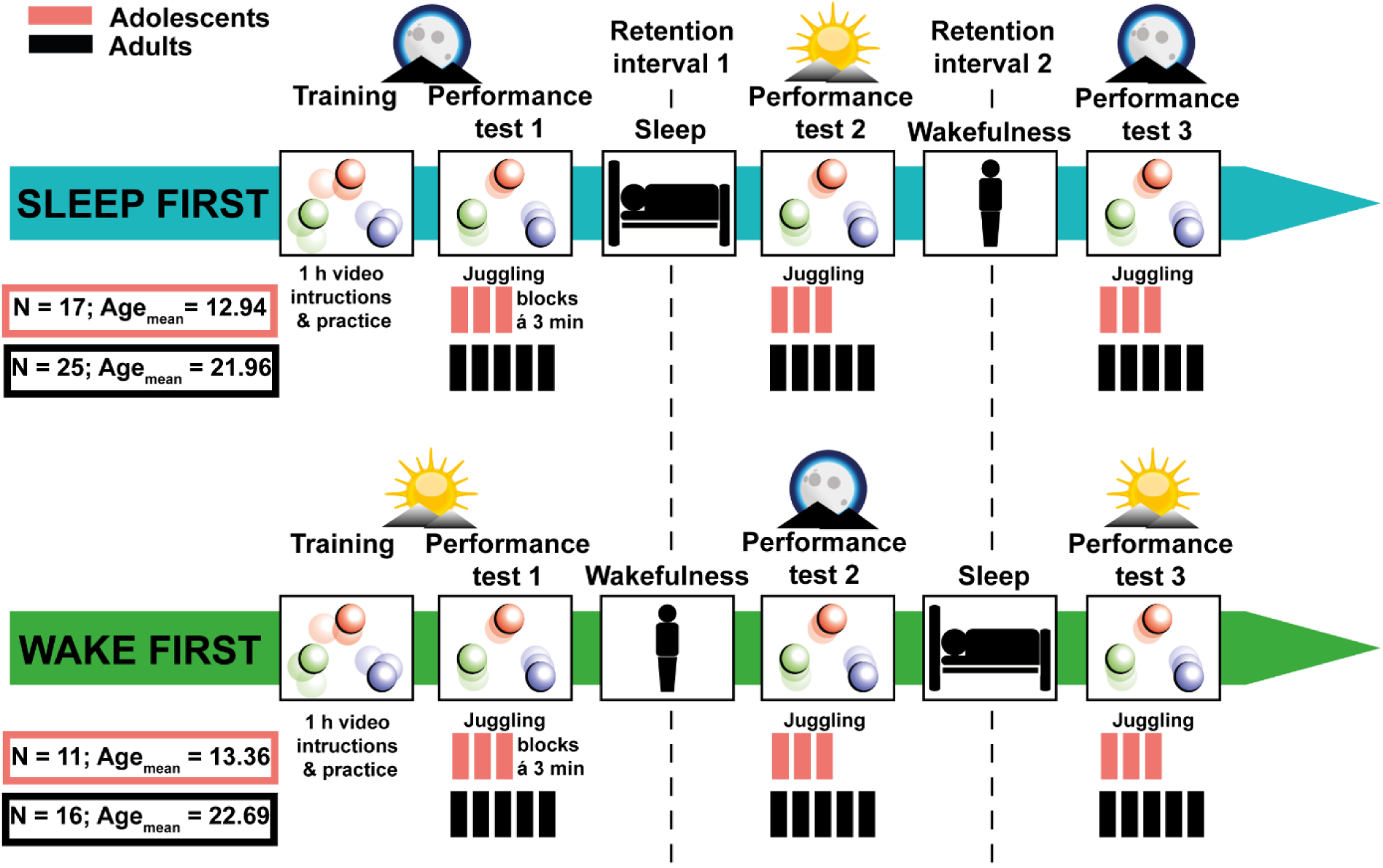
Study design. Adolescents (N = 28; 23 male) and adults (N = 41; 25 male) without prior juggling experience were divided into a *sleep-first* and a *wake-first* group. Participants in the *sleep-first* group trained to juggle for 1 hour with video instructions in the evening. Juggling performance was tested before and after a retention interval containing sleep (1), followed by a third juggling test after a retention interval containing wakefulness (2). Participants in the *wake-first* group followed the same protocol but in reverse order (i.e. training in the morning, first retention interval containing wakefulness and second retention interval containing sleep). Polysomnography during an adaptation night and a learning night at the respective sleep retention interval. Psychomotor vigilance tasks were conducted before each performance test. Adolescents only performed three juggling blocks per test to avoid a too excessive training-load.

**Figure 2.**
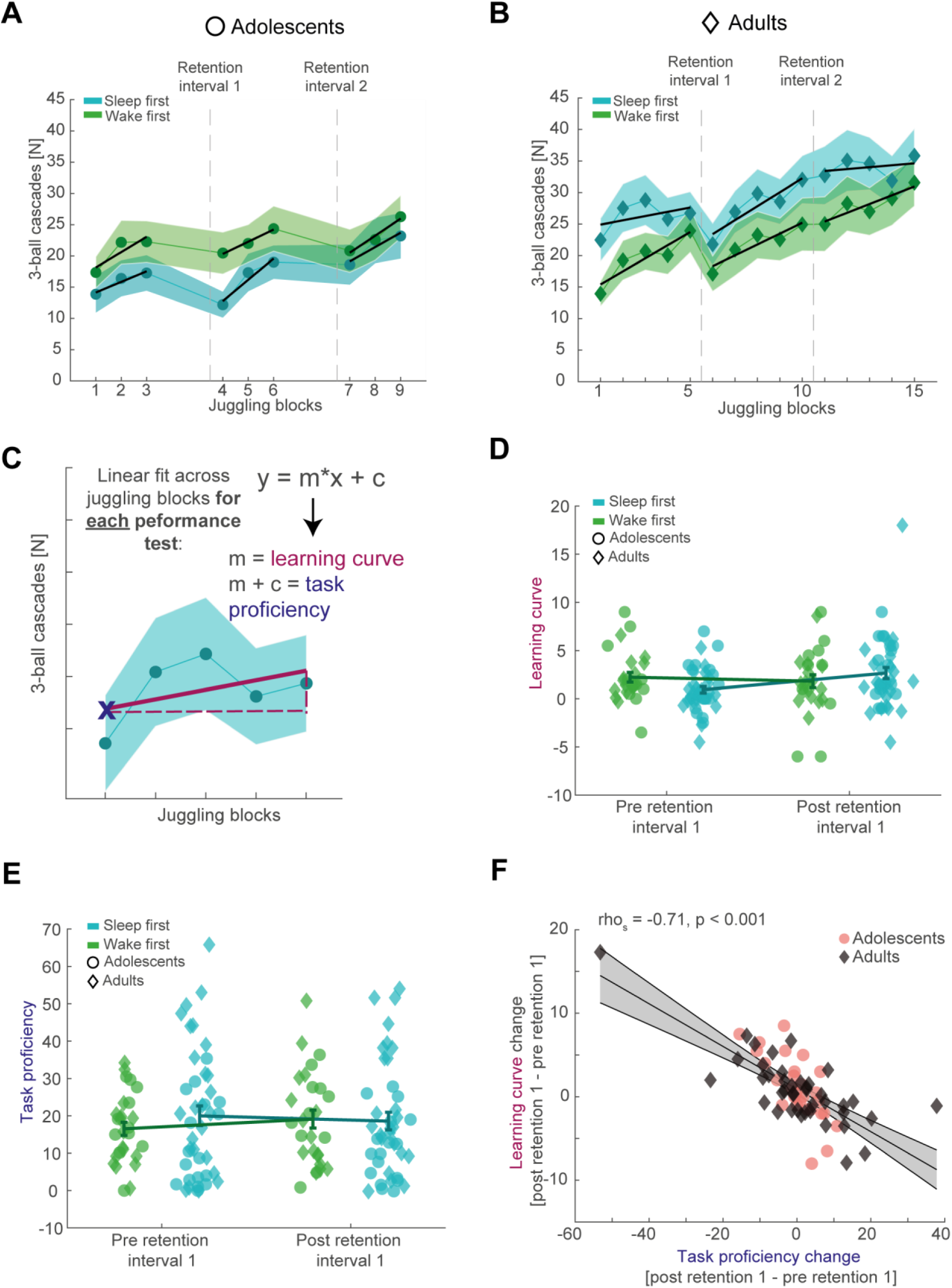
Behavioral results and parameterizing juggling performance. (**A**) Number of successful three-ball cascades (mean ± standard error of the mean [SEM]) of adolescents (circles) for the *sleep-first* (blue) and *wake-first* group (green) per juggling block. Grand average learning curve (black lines) as computed in (C) are superimposed. Dashed lines indicate the timing of the respective retention intervals that separate the three performance tests. Note that adolescents improve their juggling performance across the blocks. (**B**) Same conventions as in (A) but for adults (diamonds). Similar to adolescents, adults improve their juggling performance across the blocks regardless of group. (**C**) Schematic representation of the juggling learning process parameterization. We used a linear fit across all juggling blocks within a performance test to estimate the learning curve (m) and the task proficiency (linear line equation solved for x = 1) for each corresponding performance test. (**D**) Comparison of the juggling learning curve (mean ± SEM) between the *sleep-first* (blue) and the *wake-first* group (green) of adolescents (circles) and adults (diamonds) before and after the first retention interval to investigate the influence of sleep. Single subject data is plotted in the corresponding group color and age icon. Participants in the *sleep-first* group showed a steeper learning curve than the *wake-first* group after the first retention interval. (**E**) Same conventions as in (D) but for the task proficiency metric. Adolescents in the *wake-first* group had better overall task proficiency than adolescents in the *sleep-first* group. Adults in the *sleep-first* group displayed better overall task proficiency than adults in the *wake-first* group. (**F**) Spearman rank-correlation between the overnight change in task proficiency (post – pre retention interval) and the overnight change in learning curve with robust linear trend line collapsed over the whole sample. Grey-shaded area indicates 95% confidence intervals of the trend line. Adolescents are denoted as red circles and adults as black diamonds. A strong inverse relationship indicated that participants with an improved task proficiency show flatter learning curves.

### Behavioral results: juggling performance and disentangling the learning process

Adolescents improved their juggling performance over the course of all nine blocks (**Figure 2A top**; F_3.957, 94.962_ = 6.948, p < 0.001, η^2^ = 0.23). There was neither an overall difference in performance between the *sleep-first* and the *wake-first* group (F_1, 24_ = 1.002, p = 0.327, η^2^ = 0.04), nor did they differ over the course of the juggling blocks (F_3.957, 94.962_ = 1.148, p = 0.339, η^2^ = 0.05). Similar to the adolescents, adults improved in performance across all 15 blocks (**Figure 2B top**; F_4.673, 182.241_ = 11.967, p

< 0.001, η^2^ = 0.24), regardless of group (F_4.673, 182.241_ = 0.529, p = 0.742, η^2^ = 0.01).

Further, there was no overall difference in performance between the *sleep-first* and *wake-first* groups in adults (F_1, 39_ = 1.398, p = 0.244, η^2^ = 0.04). Collectively, these results show, that participants do not reach asymptotic level juggling performance (for single subject data of good and bad performers see **Figure 2 – figure supplement 1AB**). In other words, the gross-motor skill learning process is still in progress in adolescents and adults. Therefore, we wanted to capture the progression of the learning process, rather than absolute performance metrics (i.e. mean performance) that would underestimate the dynamics of gross-motor learning.

Since subjects did not asymptotic level performance, but learning was ongoing, we parameterized the juggling learning process by estimating the learning curve for each performance test using a first-degree polynomial fit to the different blocks (**Figure 2C; Figure 2AB**, black lines). We considered the slope of the resulting trend as learning curve. The learning process of complex motor skills is thought to consist of a fast initial learning stage during skill acquisition and a much slower skill retaining learning stage (Dayan & Cohen, 2011; Doyon & Benali, 2005). In other words, within-learning session performance gains are rapid at the beginning, but taper off with increased motor skill proficiency, resembling a power-law curve. Therefore, we also estimated the task proficiency per performance test at the first time point as predicted by the model, since the learning curve is expected to be influenced by the individual juggling aptitude. Importantly, the estimated task proficiency was comparable to the observed values in the corresponding first juggling block (performance test 1: rho_s_ = .98, p < 0.001; performance test 2: rho_s_ = .97, p < 0.001). Besides having a more accurate picture of juggling performance, this parameterization also allowed us to compare performance of adolescents and adults on a similar scale because of the different number of juggling blocks. A mixed ANOVA with the factors performance test (pre and post retention interval), condition group (*sleep-first* and *wake-first*) and age group (adolescents and adults) showed a significant interaction between performance test and condition group (F_1, 65_ = 4.868, p = 0.031, η^2^ = 0.07). This result indicates that regardless of age, the juggling learning curve becomes steeper after sleep than after wakefulness, thus indicating that sleep impacts motor learning (**Figure 2D**). No other interactions or main effects were significant (for the complete ANOVA report see **Supplementary file - table 3**). When analyzing the task proficiency before and after the first retention interval, depending on condition and age group, we found a significant interaction between condition and age group (**Figure 2E**; F_1, 65_ = 5.210, p = 0.026, η^2^ = 0.07), showing that the adult *sleep-first* group had better overall task proficiency than the *wake-first* group, whereas the adolescent *sleep-first* group was worse than the *wake-first* group. The interaction (performance test x condition group) did not reach significance (F_1, 65_ = 1.882, p = 0.175, η^2^ = 0.03; also see **Supplementary file - table 4**). Collectively, these results suggest that sleep influences learning of juggling as a gross motor task. **Figure 2A** and **2B** indicate that performance tests in the morning might be characterized by a steeper learning curve than the evening tests. We confirmed this observation using a linear mixed model (**Supplementary file – table 5AB**). While this finding might also indicate a circadian influence on learning in our task, we did not find evidence for an effect on circadian sensitive psychomotor vigilance task reaction time. Neither when comparing sleep first and wake first groups (**Figure 2 – figure supplement 1C**), nor when specifically probing evening and morning performance tests (**Supplementary file – table 5EF**).

Next, we further dissected the relationship between changes in the learning curve and task proficiency after the retention interval. We hypothesized, that a stronger increase in task proficiency across sleep would lead to a flatter learning curve based on the assumption that motor skill learning involves fast and slow learning stages. Indeed, we confirmed a strong negative correlation between the change (post retention values – pre retention values) in task proficiency and the change in learning curve after the retention interval (**Figure 2F**; rho_s_ = -0.71, p < 0.001), which also remained strong after outlier removal (**Figure 2 – figure supplement 1D**). This result indicates that participants who consolidate their juggling performance after a retention interval show slower gains in performance. Note, that the flattening of the learning curve does not necessarily indicate worse learning but rather mark a more progressed learning stage. These results demonstrate a highly dynamic gross-motor skill learning process. Given that sleep influences the juggling learning curve, we aimed to determine whether sleep oscillation dynamics track the dynamics of gross-motor learning.

### Electrophysiological results: inter-individual variability and SO-spindle coupling

To determine the nature of the timed coordination between the two cardinal sleep oscillations, we adopted the same principled individualized approach we developed earlier (Hahn et al., 2020). First, we compared oscillatory power between adolescents and adults in the frequency range between 0.1 and 20 Hz during NREM (2&3) sleep, using cluster-based permutation tests (Maris & Oostenveld, 2007). Spectral power was elevated in adolescents as compared to adults across the whole tested frequency range (**Figure 3 – figure supplement 1A left** for representative electrode Cz; cluster test: p < 0.001, d = 1.88). Similar to the previously reported developmental patterns of sleep oscillations from childhood to adolescence (Hahn et al., 2020), this difference was explained by a spindle frequency peak shift and broadband decrease in the fractal or 1/f trend of the signal, thus directly replicating and extending our previous findings in a separate sample. After estimating the fractal component of the power spectrum by means of irregular-resampling auto-spectral analysis (Wen & Liu, 2016), we found that adolescents exhibited a higher offset of fractal component on the y-axis than adults (**Figure 3 – figure supplement 1A middle**; cluster test: p < 0.001, d = 1.99). Next, we subtracted the fractal component from the power spectrum, which revealed clear distinct oscillatory peaks in the SO (< 2 Hz) and sleep spindle range (11 – 16 Hz) for both, adolescents and adults (**Figure 3 – figure supplement 1A right**). Importantly, we observed the expected spatial amplitude topography with stronger frontal SO and pronounced centro-parietal spindles for both age groups (**Figure 3A left**).

**Figure 3.**
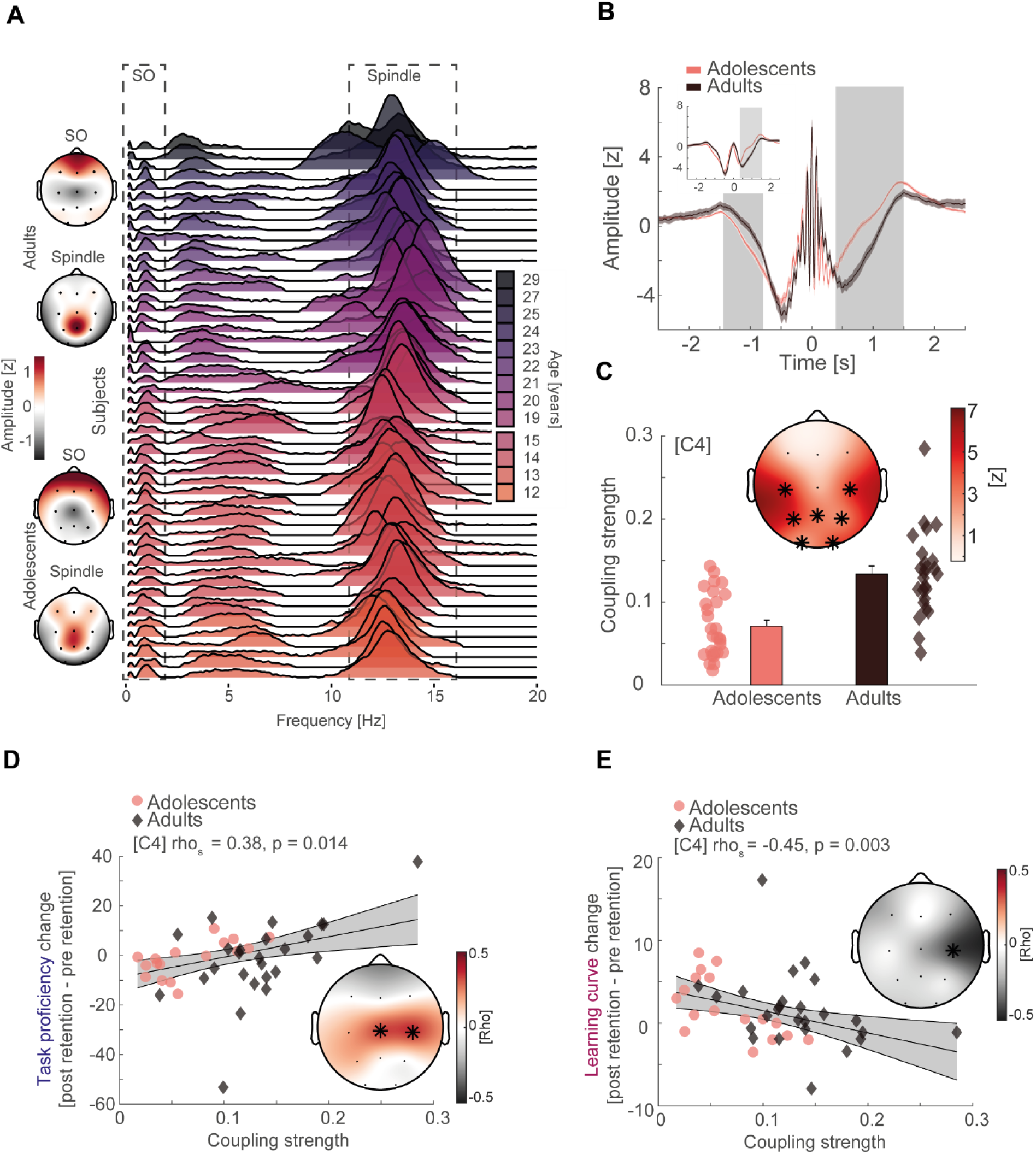
Inter-individual variability, SO-spindle coupling development, and neural correlates of gross-motor learning dynamics. (**A**) Left: topographical distribution of the 1/f corrected SO and spindle amplitude as extracted from the oscillatory residual (Figure 3 – figure supplement 1A, right). Note that adolescents and adults both display the expected topographical distribution of more pronounced frontal SO and centro-parietal spindles. Right: single subject data of the oscillatory residual for all subjects with sleep data color coded by age (darker colors indicate older subjects). SO and spindle frequency ranges are indicated by the dashed boxes. Importantly, subjects displayed high inter-individual variability in the sleep spindle range and a gradual spindle frequency increase by age that is critically underestimated by the group average of the oscillatory residuals (Figure 3 – figure supplement 1A, right). (**B**) Spindle peak locked epoch (NREM3, co-occurrence corrected) grand averages (mean ± SEM) for adolescents (red) and adults (black). Inset depicts the corresponding SO-filtered (2 Hz lowpass) signal. Grey-shaded areas indicate significant clusters. Note, we found no difference in amplitude after normalization. Significant differences are due to more precise SO-spindle coupling in adults. (**C**) Top: comparison of SO- spindle coupling strength between adolescents and adults. Adults displayed more precise coupling than adolescents in a centro-parietal cluster. T-scores are transformed to z-scores. Asterisks denote cluster-corrected two-sided p < 0.05. Bottom: Exemplary depiction of coupling strength (mean ± SEM) for adolescents (red) and adults (black) with single subject data points. Exemplary single electrode data (bottom) is shown for C4 instead of Cz to visualize the difference. (**D**) Cluster-corrected correlations between individual coupling strength and overnight task proficiency change (post – pre retention) for adolescents (red, circle) and adults (black, diamond) of the sleep-first group (left, data at C4). Asterisks indicate cluster-corrected two-sided p < 0.05. Grey-shaded area indicates 95% confidence intervals of the trend line. Participants with a more precise SO-spindle coordination show improved task proficiency after sleep. Note that the change in task proficiency was inversely related to the change in learning curve (cf. Figure 2D), indicating that a stronger improvement in task proficiency related to a flattening of the learning curve. Further note that the significant cluster formed over electrodes close to motor areas. (**E**) Cluster-corrected correlations between individual coupling strength and overnight learning curve change. Same conventions as in (D). Participants with more precise SO-spindle coupling over C4 showed attenuated learning curves after sleep.

Critically, the displayed group averages of the oscillatory residuals (**Figure 3 – figure supplement 1A right**) underestimate the inter-individual variability of the spindle frequency peak (**Figure 3A right**; oscillatory residuals for all subjects at Cz). Even though we found the expected systematic spindle frequency increase in a fronto- parietal cluster from adolescence to adulthood (**Figure 3 – figure supplement 1B**; cluster test: p = 0.002, d = -0.87), both respective age groups showed a high degree of variability of the inter-individual spindle peak.

Based on these findings, we separated the oscillatory activity from the fractal activity for every subject at every electrode position to capture the individual features of SO and sleep spindle oscillations. We then used the extracted individual features from the oscillatory residuals to adjust SO and spindle detection algorithms (Hahn et al., 2020; Helfrich et al., 2018; Molle et al., 2011; Staresina et al., 2015) to account for the spindle frequency peak shift and high inter-individual variability. To ensure the simultaneous presence of the two interacting sleep oscillations in the signal, we followed a conservative approach and restricted our analyses to NREM3 sleep given the low co-occurrence rate in NREM2 sleep (**Figure 3 – figure supplement 1CD**) which can cause spurious coupling estimates (Hahn et al., 2020). Further, we only considered spindle events that displayed a concomitant detected SO within a 2.5 s time window.

We identified an underlying SO component (2 Hz low-pass filtered trace) in the spindle peak locked averages for adolescents and adults on single subject and group average basis (**Figure 3 – figure supplement 1E**), indicating a temporally precise interaction between sleep spindles and SO that is clearly discernible in the time domain.

To further assess the interaction between SO and sleep spindles, we computed SO-trough-locked time-frequency representations (**Figure 3 – figure supplement 1F**). Adolescents and adults revealed a shifting temporal pattern in spindle activity (11 – 16 Hz) depending on the SO phase. In more detail, spindle activity decreased during the negative peak of the SO (‘down-state’) but increased during the positive peak (‘up- state’). This temporal pattern and the underlying SO-component in spindle event detection (**Figure 3 – figure supplement 1E**) confirm the coordinated nature of the two major sleep oscillations in adolescents and adults.

Next, we determined the coordinated interplay between SO and spindles in more detail by analyzing individualized event-locked cross-frequency interactions (Dvorak & Fenton, 2014; Hahn et al., 2020; Helfrich et al., 2019). In brief, we extracted the instantaneous phase angle of the SO-component (< 2 Hz) corresponding to the positive spindle amplitude peak for all trials at every electrode per subject. We assessed the cross frequency coupling based on z-normalized spindle epochs (**Figure 3B**) to alleviate potential power differences due to age (**Figure 3 – figure supplement 1A)** or different EEG-amplifier systems that could potentially confound our analyses (Aru et al., 2015). Importantly, we found no amplitude differences around the spindle peak (point of SO-phase readout) between adolescents and adults using cluster-based random permutation testing (**Figure 3B**), indicating an unbiased analytical signal. This was also the case for the SO-filtered (< 2 Hz) signal (**Figure 3B**, inset). Critically, the significant differences in amplitude from -1.4 to -0.8 s (p = 0.023, d = -0.73) and 0.4 to 1.5 s (p < 0.001, d = 1.1) are not caused by age related differences in power or different EEG-systems but instead by the increased coupling strength (i.e. higher coupling precision of spindles to SOs) in adults giving rise to a more pronounced SO-wave shape when averaging across spindle peak locked epochs. Further, we specifically focused our analyses on spindle events to account for the higher variability in the spindle frequency band than in the SO-band (**Figure 3A**). Based on these adjusted phase values, we derived the coupling strength defined as 1 – circular variance. This metric describes the consistency of the SO-spindle coupling (i.e. higher coupling strength indicates more precise coupling) and has previously been shown to accurately track brain development and memory formation (Hahn et al., 2020). As expected, adults had a higher coupling strength in a centro-parietal cluster than adolescents (**Figure 3C**; cluster test: p < 0.001, d = 0.88), indicating a more precise interplay between SO and spindles during adulthood.

### SO-spindle coupling tracks gross-motor learning

After demonstrating that SO-spindle coupling becomes more precise from early adolescence to adulthood, we tested the hypothesis, that the dynamic interaction between the two sleep oscillations explains the dynamic process of complex gross-motor learning. When taking the behavioral analyses into account, we did not find any evidence for a difference between the two age groups on the impact of sleep on the learning curve (**Figure 2D**). Therefore, we did not differentiate between adolescents and adults in our correlational analyses. Furthermore, given that we only recorded polysomnography for the adults in the sleep first group and that adolescents in the wake first group showed enhanced task proficiency at the time point of the sleep retention interval due to additional training (**Figure 3 – figure supplement 2A**), we only considered adolescents and adults of the *sleep-first* group to ensure a similar level of juggling experience (for summary statistics of sleep architecture and SO and spindle events of subjects that entered the correlational analyses see **Supplementary file – table 6**). Notably, we found no differences in electrophysiological parameters (i.e. coupling strength, event detection) between the adolescents of the wake first and sleep first group (**Figure 3 – figure supplement 2B** & **Supplementary file – table 7**). To investigate whether coupling strength in the night of the first retention interval explains overnight changes of task proficiency (post retention interval 1 – pre retention interval 1), we computed cluster-corrected correlation analyses. We identified a significant central cluster (**Figure 3D**; mean rho = 0.37, p = 0.017), indicating that participants with a more consistent SO-spindle interplay have stronger overnight improvements in task proficiency.

Given that we observed a strong negative correlation between task proficiency at a given time point and the steepness of the subsequent learning curve (cf. **Figure 2F**) as subjects improve but do not reach ceiling level performance, we conversely expected a negative correlation between learning curve and coupling. Given this dependency, we observed a significant cluster-corrected correlation at C4 (**Figure 3E**; rho_s_ = -0.45, p = 0.039, cluster-corrected), showing that participants with a more precise SO-spindle coupling exhibit a flatter learning curve overnight. This observation is in line with a trade-off between proficiency and learning curve, which exhibits an upper boundary (100% task proficiency). In other words, individuals with high performance exhibit a smaller gain through additional training when approaching full task proficiency.

Critically, when computing the correlational analyses separately for adolescents and adults, we identified highly similar effects at electrode C4 for task proficiency (**Figure 3 – figure supplement 2C**) and learning curve (**Figure 3 – figure supplement 2D**) in each group. These complementary results demonstrate that coupling strength predicts gross-motor learning dynamics in both, adolescents as well as adults, and further shows that this effect is not solely driven by one group. Furthermore, our results remained consistent when including coupled spindle events in NREM2 (**Figure 3 – figure supplement 2E**) and after outlier removal (**Figure 3 – figure supplement 2FG**).

To rule out age as a confounding factor that could drive the relationship between coupling strength, learning curve and task proficiency in the mixed sample, we used cluster-corrected partial correlations to confirm their independence of age differences (task proficiency: mean rho = 0.40, p = 0.017; learning curve: rho_s_ = -0.47, p = 0.049). Additionally, given that we found that juggling performance could underlie a circadian modulation we controlled for individual differences in alertness between subjects due to having just slept. We partialed out the mean PVT reaction time before the juggling performance test after sleep from the original analyses and found that our results remained unchanged (task proficiency: mean rho = 0.37, p = 0.025; learning curve: rho_s_ = -0.49, p = 0.040). For a summary of the reported cluster-corrected partial correlations as well as analyses controlling for differences in sleep architecture see **Figure 3 – figure supplement 3**. Further, we also confirmed that our correlations are not influenced by individual differences in SO and spindle event parameters (**Figure 3 – figure supplement 4).**

Finally, we investigated whether subjects with high coupling strength have a gross-motor learning advantage (i.e. trait-effect) or a learning induced enhancement of coupling strength is indicative for improved overnight memory change (i.e. state-effect). First, we correlated SO-spindle coupling strength obtained from the adaptation night with the coupling strength in the learning night. We found that overall, coupling strength is highly correlated between the two measurements (mean rho across all channels = 0.55, **Figure 3 – figure supplement 2H**), supporting the notion that coupling strength remains rather stable within the individual (i.e. trait). Second, we calculated the difference in coupling strength between the learning night and the adaptation night to investigate a possible state-effect. We found no significant cluster-corrected correlations between coupling strength change and task proficiency- as well as learning curve change (**Figure 3 – figure supplement 2I**).

Collectively, these results indicate the regionally specific SO-spindle coupling over central EEG sensors encompassing sensorimotor areas precisely indexes learning of a challenging motor task.

## DISCUSSION

By comparing adolescents and adults learning a complex juggling task, we critically advance our previous work about the intricate interplay of learning and memory formation, brain maturation and coupled sleep oscillations: First, we demonstrated that SO-spindle interplay precision is not only enhanced from childhood to late adolescence but also progressively improves from early adolescence to young adulthood (**Figure 3F**). Second and more importantly, we provide first evidence that the consistency of SO-spindle coordination is a promising model to track real-life gross-motor skill learning in addition to its key role in declarative learning (**Figure 3DE**). Notably, this relationship between coupling and learning occurred in a regional specific manner and was pronounced over frontal areas for declarative and over motor regions for procedural learning (Hahn et al., 2020). Collectively, our results suggest that precise SO-spindle coupling supports gross-motor memory formation by integrating information from subcortical memory structures to cortical networks.

How do SO-spindle interactions subserve motor memory formation? Motor learning is a process relying on complex spatial and temporal scales in the human brain. To acquire motor skills the brain integrates information from extracortical structures with cortical structures via cortico-striato-thalamo-cortico loops and cortico-cerebello-thalamo-cortico circuits (Dayan & Cohen, 2011; Doyon & Benali, 2005; Doyon et al., 2018; Pinsard et al., 2019). However, growing evidence also advocates for hippocampal recruitment for motor learning, especially in the context of sleep-dependent memory consolidation (Albouy et al., 2013; Boyke et al., 2008; Draganski et al., 2004; Pinsard et al., 2019; Sawangjit et al., 2018; Schapiro et al., 2019). Hippocampal memory reactivation during sleep is one cornerstone of the active systems consolidation theory, where coordinated SO-spindle activity route subcortical information to the cortex for long-term storage (Diekelmann & Born, 2010; Helfrich et al., 2019; Klinzing et al., 2019; Ngo et al., 2020). Quantitative markers of spindle and SO activity but not the quality of their interaction have been frequently related to motor memory in the past (Barakat et al., 2011; Bothe et al., 2019; Bothe et al., 2020; Huber et al., 2004; Morita et al., 2012; Nishida & Walker, 2007; Tamaki et al., 2008). Our results now complement the active systems consolidation theories’ mechanistic assumption of interacting oscillations by demonstrating that a precise SO-spindle interplay subserves gross-motor skill learning (**Figure 3DE**). Of note, we did not derive direct hippocampal activity in the present study given spatial resolution of scalp EEG-recordings. Nonetheless, as demonstrated recently, coupled spindles precisely capture cortico-hippocampal network communication as well as hippocampal ripple expression (Helfrich et al., 2019). Thus, higher SO-spindle coupling strength supporting gross-motor learning in our study points towards a more efficient information exchange between hippocampus and cortical areas.

Remarkably, hippocampal engagement is especially crucial at the earlier learning stages. Recently, it has been found that untrained motor sequences exhibit hippocampal activation that subsides for more consolidated sequences. This change was further accompanied by increased motor cortex activation, suggesting a transformative function of sleep for motor memory (Pinsard et al., 2019). In other words, hippocampal disengagement likely indexes the transition from the fast learning stage to the slower learning stage with more proficient motor skill (Dayan & Cohen, 2011; Doyon & Benali, 2005). The dynamics of the two interacting learning stages of motor skill acquisition are likely reflected by the inverse relationship between task proficiency increases and learning curve attenuation (**Figure 2F**). Given that our subjects did not reach asymptotic performance level (**Figure 2AB**) and that SO-spindle coupling tracks gross-motor skill learning dynamics as it relates to both, learning curve attenuation and task proficiency increments, it is plausible that SO-coupling strength represents the extent of hippocampal support for integrating information to motor cortices during complex motor skill learning.

Interestingly, SO and spindles are not only implicated in hippocampal-neocortical network communication but are also indicative for activity and information exchange in subcortical areas that are more traditionally related to the shift from fast to slow motor learning stages. For example, striatal network reactivation during sleep was found to be synchronized to sleep spindles, which predicted motor memory consolidation (Fogel et al., 2017). In primates, coherence between M1 and cerebellum in the SO and spindle frequency range suggested that coupled oscillatory activity conveys information through cortico-thalamo-cerebellar networks (Xu et al., 2020). One testable hypothesis for future research is whether SO-spindle coupling represents a more general gateway for the brain to exchange subcortical and cortical information and not just hippocampal-neocortical communication.

Critically, we found that the consistency of the SO-spindle interplay identified at electrodes overlapping with motor areas such as M1 was predictive for the gross-motor learning process (**Figure 3DE**). This finding corroborates the idea that SO-spindle coupling supports the information flow between task-relevant subcortical and cortical areas. Recent evidence in the rodent model demonstrated that neural firing patterns in M1 during spindles became more coherent after performing a grasping motor task. The extent of neural firing precision was further mediated by a function of temporal proximity of spindles to SOs (Silversmith et al., 2020). Through this synchronizing process and their Ca2+ influx propagating property, coupled spindles are likely to induce neural plasticity that benefits motor learning (Niethard et al., 2018).

How relevant is sleep for real-life gross-motor memory consolidation? We found that sleep impacts the learning curve but did not affect task proficiency in comparison to a wake retention interval (**Figure 2DE**). Two accounts might explain the absence of a sleep effect on task proficiency. (1) Sleep rather stabilizes than improves gross-motor memory, which is in line with previous gross-motor adaption studies (Bothe et al., 2019; Bothe et al., 2020). (2) Pre-sleep performance is critical for sleep to improve motor skills (Wilhelm et al., 2012). Participants commonly reach asymptotic pre-sleep performance levels in finger tapping tasks, which is most frequently used to probe sleep effects on motor memory. Here we found that using a complex juggling task, participants do not reach asymptotic ceiling performance levels in such a short time. Indeed, the learning progression for the *sleep-first* and *wake-first* groups followed a similar trend (**Figure 2AB**), suggesting that more training and not in particular sleep drove performance gains. We note that juggling performance in our study could have been influenced by the timing of when learning is optimal in the circadian cycle. However, we did not find evidence for a circadian modulation of cognitive engagement based on objective reaction time data (**Figure 2 – figure supplement 1C**). Nonetheless, we cannot fully disentangle circadian and sleep effects with our study design, which should be considered a limitation to our findings. Importantly, SO-spindle coupling still predicted learning dynamics on a single subject level advocating for a supportive function of sleep for gross-motor memory. Moreover, we found that SO-spindle coupling strength remains remarkably stable between two nights, which also explains why a learning-induced change in coupling strength did not relate to behavior (**Figure 3 – figure supplement 2I**). Thus, our results primarily suggest that strength of SO-spindle coupling correlates with the ability to learn (trait), but does not solely convey the recently learned information. This set of findings is in line with recent ideas that strong coupling indexes individuals with highly efficient subcortical-cortical network communication (Helfrich et al., 2021).

This subcortical-cortical network communication is likely to be refined throughout brain development, since we discovered elevated coupling strength in adults compared to early adolescents (**Figure 3C**). This result compliments our earlier findings of enhanced coupling precision from childhood to adolescence (Hahn et al., 2020) and the recently demonstrated lower coupling strength in pre-school children (Joechner et al., 2021). We speculate that, similar to other spindle features, the trajectory of SO-coupling strength is likely to reach a plateau during adulthood (Nicolas et al., 2001; Purcell et al., 2017). Importantly, we identified similar methodological challenges to assess valid cross-frequency coupling estimates in the current cross-sectional study to the previous longitudinal study. Age severely influences fractal dynamics in the brain (**Figure 3 – figure supplement 1A**) and the defining features of sleep oscillations (**Figure 3B & Figure 3 – figure supplement 1B**). Remarkably, inter-individual oscillatory variability was pronounced even in the adult age group (**Figure 3A**), highlighting the critical need to employ individualized cross-frequency coupling analyses to avoid its pitfalls (Aru et al., 2015; Muehlroth & Werkle-Bergner, 2020).

Taken together, our results provide a mechanistic understanding of how the brain forms real-life gross-motor memory during sleep. As sleep has been shown to support fine-motor memory consolidation in individuals after stroke (Gudberg & Johansen-Berg, 2015; Siengsuhon & Boyd, 2008), SO-spindle coupling integrity could be a valuable, easy to assess predictive index for rehabilitation success.

## ACKNOWLEDGMENTS

This research was supported by Austrian Science Fund (P25000-B24) and the Centre for Cognitive Neuroscience Salzburg (CCNS). M.A.H. was additionally supported by the Doctoral Collage ‘Imaging the Mind’ (FWF, Austrian Science Fund W1233-G17). R.F.H. is supported by the German Research Foundation (DFG, HE 8329/2-1), the Hertie Foundation (Hertie Network of Excellence in Clinical Neurosciences) and the Jung Foundation for Science and Research (Ernst Jung Career Advancement Award).

## AUTHOR CONTRIBUTIONS

Conceptualization, M.A.H., R.F.H., K.H.; Methodology, R.F.H., K.H., M.S.; Software, R.F.H., M.A.H.; Validation, M.A.H., R.F.H., K.H.; Formal Analysis, M.A.H.; Investigation, M.A.H., K.B., K.H.; Resources, K.H., M.S.; Data Curation, M.A.H., K.H., K.B., D.P.J.H.; Writing – Original Draft, M.A.H.; Writing – Review & Editing, R.F.H., K.H., M.S., D.P.J.H., K.B.; Visualization, M.A.H., D.P.J.H; Supervision, R.F.H, K.H.; Project Administration, K.H.; Funding Acquisition, K.H.

## MATERIAL AND METHODS

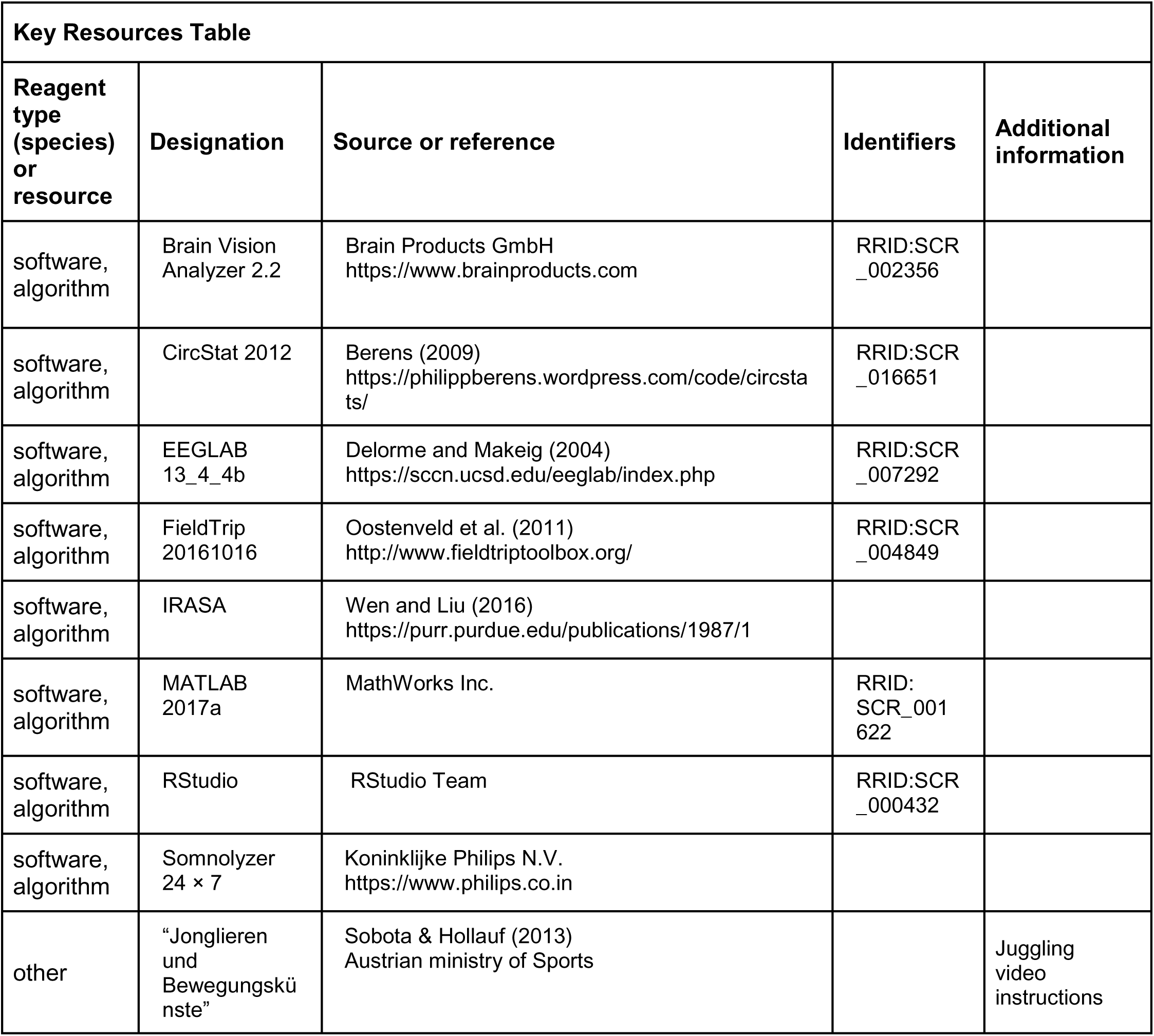

### Participants

We recruited 29 adolescents (mean ± SD age, 13.17 ± 0.85 years; 5 female, 24 male) from a local boarding school and 41 young adults (mean ± SD age, 22.24 ± 2.15 years; 16 female, 25 male) from the student population of the University of Salzburg. All participants were healthy, right-handed and without prior juggling experience. However, we excluded one adolescent for all analyses post-hoc for violating the prior juggling experience criteria. Two adolescents did not participate in the third performance test. We randomly divided adolescents and adults into a *sleep-first* (adolescents: N = 17, 12.94 ± 0.75 years; 3 females, 14 males; adults: N = 25, 21.95 ± 2.42 years; 8 females, 17 males) and a *wake-first* group (adolescents: N = 11, 13.36 ± 0.81 years; 2 females, 9 males; adults: N = 16, 22.69 ± 1.62 years; 8 females, 8 males). See experimental design for more detailed information about the groups. We recorded polysomnography (PSG) during full night sleep for all participants except adults in the *wake-first* group. Therefore, comparison of electrophysiological data between adults and adolescents was based on the adult *sleep-first* group and both adolescent groups. To ensure similar juggling learning experience, we only included adults and adolescents in the *sleep-first* group when analyzing the relationship between electrophysiological measures and behavioral performance. All participants and the legal custodians of the adolescents provided written informed consent before participating in the study. The study protocol was conducted in accordance with the Declaration of Helsinki and approved by the ethics committee of the University of Salzburg (EK-GZ:16/2014). Adults received monetary compensation or student credit for their participation. Adolescents received a set of juggling balls.

### Experimental design

Adults in the *sleep-first* group visited the sleep laboratory on three occasions (**Figure 1**). At the first day subjects slept in the sleep lab with full night PSG for adaptation purposes. On the second visit, subjects learned and practiced juggling by video instructions in the evening (8.45 pm - 9.45 pm). Juggling performance was assessed three times in total. The first performance test was conducted after the training session (10.00 pm – 10.18pm). The second performance test (7.30 am – 7.48 am) took place after the first retention interval containing a full night of sleep with polysomnography (11 pm – 7 am). The third and last performance test was executed after the second retention interval (9.00 pm – 9.18 pm) containing wakefulness. Adults in the *wake-first* group followed a similar protocol but with reversed order of the retention intervals (i.e. first retention interval containing wakefulness and the second interval containing sleep). Therefore, participants performed the juggling training (10.15 am – 11.15 am) and the first performance test (11.30 am – 11.48 am) in the morning, the second performance test after wakefulness (9.00 pm – 9.18 pm) and the third performance test after sleep (11.00 am – 11.18 am). We did not record polysomnography in the *wake-first* group because participants slept at home. To objectively assess attentiveness and potential circadian influences, all participants completed a psychomotor vigilance task (Dinges & Powell, 1985) before the performance tests. Actigraphy (Cambridge Neurotechnology Actiwatch, Cambridge, UK) and a sleep log (Saletu et al., 1987) verified compliance with a regular sleep schedule throughout the study.

Adolescents went through a study protocol comparable to the adults. However, we adjusted the protocol to adhere to the schedule of the boarding school and to control the training load. First, we recorded ambulatory PSG for both groups in their habitual sleep environment at the boarding school and second, we reduced the number of juggling blocks during the performance tests (for details see gross-motor task) because the study regime was already exhausting for our adult participants and we wanted to avoid a too excessive training load. The *sleep-first* group performed the juggling training (6.30 pm – 7.30 pm) and performance test in the evening (7.45 pm – 7.58 pm) followed by a retention interval containing sleep (21.00 pm – 6.00 am). The second performance test was conducted after sleep (7.30 am – 7.43 am) and the third performance test after wakefulness (7.30 pm – 7.43 pm). The *wake-first* group learned to juggle (7.30 am – 8.30 am) with a subsequent performance test (8.45 am – 8.58 am) in the morning. The second performance test was executed after wakefulness in the evening (7.30 pm – 7.43 pm) and the third performance test was completed after sleep (7.30 am – 7.43 am).

### Gross-motor task

To investigate the involvement of slow oscillation-spindle coupling in acquiring a real-life gross motor skill, we implemented a juggling paradigm, which has been shown to induce neural plasticity (Boyke et al., 2008; Draganski et al., 2004) and to be sensitive for sleep-dependent memory consolidation (Morita et al., 2012, 2016). Adults and adolescents completed the same juggling training, which was based on short video clips from the “Juggling and Movement Arts” DVD (“Jonglieren und Bewegungskünste”; Sobota & Hollauf, 2013) containing step-by-step instructions from the correct stance to a full five-ball cascade (i.e. five continuous catches). We used 14 video clips demonstrating the exercises followed by a practice opportunity for the participants. The training session lasted approximately one hour with a short break after half an hour. During the performance tests, participants were instructed to juggle as accurately and continuously as possible. Adults juggled for five blocks á three minutes, which was always separated by a 30 second break. To alleviate the physical strain, adolescents only juggled for three blocks á three minutes during the performance tests. Training and performance tests were videotaped to evaluate the juggling performance.

### Parameterizing juggling performance

We evaluated the juggling performance by counting consecutive catches based on the video material. We used the number of three ball cascades (i.e. three catches in a row, **Figure 2AB**) as index for juggling performance by dividing the number of consecutive catches by three. We opted for three ball cascades as a performance index because we considered three consecutive catches as the criteria for the motor task to qualify as juggling (Boyke et al., 2008; Draganski et al., 2004). Because juggling is a complex motor task where it is unlikely to reach ceiling level performance, we were interested in the progression of the learning process and how it is influenced by task proficiency. Therefore, we calculated a first degree polynomial fit using the least-squares method to parameterize the learning curve (m, slope) per performance test block (**Figure 2AB**, black lines & **Figure 2CD**), using the formula:

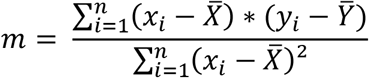

Next, we calculated the intercept c according to the following formula:

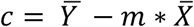

Finally, task proficiency (y_1_, **Figure 2E**) was estimated at the first time point of each performance test as

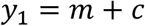

### Polysomnography and sleep staging

We recorded PSG with two systems. We conducted the ambulatory sleep recordings of the adolescents with a portable amplifier system (Alphatrace, Becker Meditec, Karlsruhe, Germany) with a sampling rate of 512 Hz. For in lab recordings of the adult participants, we utilized a 32-channel Neuroscan amplifier system (Scan 4.3.3 Software, Neuroscan Inc., Charlotte, NC) with a sampling rate of 500 Hz. Electrode placement was identical between the two recording systems and in accordance with the 10-20 system. Signals were recorded with gold cup electrodes placed at F3, Fz, F4, C3, Cz, C4, P3, Pz, P4, O1 and O2 on the scalp, as well as at A1 and A2 placed at the mastoids. To allow for sleep staging and to control for muscle artifacts, we recorded an electromyogram (EMG, bipolar electrodes at the musculus mentalis), a horizontal electrooculogram (EOG, above the right outer canthus and below the left outer canthus) and a vertical EOG (above and below the left eye). We used Cz as online reference and AFz as ground electrode. For sleep staging, we re-referenced the signal offline against contralateral mastoids. Sleep was semi-automatically staged in 30 s epochs using the Somnolyzer 24x7 algorithm (Koninklijke Philips N.V.; Eindhoven, The Netherlands) and subsequently controlled by an expert scorer according to standard sleep staging criteria (Iber et al., 2007). For all other data analyses, we demeaned and re-referenced the EEG signal to a common average.

### Individualized cross-frequency coupling

To assess the precise interplay between SO and spindles, we used the same individualized cross-frequency coupling pipeline we developed earlier in order to account for network changes induced by aging, that are known to cause spurious coupling estimates (Aru et al., 2015; Cole & Voytek, 2017; Hahn et al., 2020; Scheffer-Teixeira & Tort, 2016). In brief, our approach was based on the following principles: (1) establishing the presence of sleep oscillations, (2) individually detecting transient oscillatory events, (3) alleviating power differences and (4) ensuring co-occurrence of SO (phase providing signal) and sleep spindles (amplitude providing signal).

### Establishing sleep oscillations

First, we z-normalized the EEG-signal in the time domain to mitigate prominent power differences and computed averaged power spectra from 0.1 to 30 Hz using a Fast Fourier Transform (FFT) routine with a Hanning window on 15 s of continuous NREM sleep (i.e. NREM2 and NREM3, **Figure 3 – figure supplement 1A, left**) with a 1 s sliding window. Data are presented in the semi-log space. Next, we sought to isolate the oscillatory activity in the normalized data by means of irregular auto-spectral analysis (IRASA, (Wen & Liu, 2016)). We first derived the 1/f fractal component (**Figure 3 – figure supplement 1A middle**) from 15 s NREM sleep data in 1 s sliding steps and subsequently subtracted it from the power spectrum (**Figure 3 – figure supplement 1A left**) to obtain an unbiased estimate of the oscillatory activity for every subject on every electrode (**Figure 3 – figure supplement 1A right & Figure 3A**). To separate the 1/f component from the power spectrum, we used the same parameters as specified previously (Hahn et al., 2020). In short, the signal is stretched and compressed by the same non-integer factor (e.g. stretching by a factor of 1.1 and compressing by a factor of 0.9). We repeated the resampling with factors from 1.1 to 1.9 in 0.05 steps. This pair wise stretching and compressing systematically causes frequency peak shifts in the regular oscillatory activity but leaves the more random 1/f background activity unaffected. Because the oscillatory activity becomes faster by a similar factor as it becomes slower, the oscillatory activity is averaged out by median averaging across all pair wise resampled segments thus extracting the 1/f component. We then detected individual SO (< 2 Hz) and spindle peak frequencies (10 – 17 Hz, **Figure 3 – figure supplement 1B**) and the corresponding 1/f corrected amplitude (**Figure 3A left**) in the oscillatory residual (**Figure 3 – figure supplement 1A right**). We considered the highest peak within the specified SO and spindle frequency ranges above as the most representative oscillatory event in each electrode. We then utilized the individual frequency peaks to inform the algorithms for discrete SO and spindle event detection.

### Individually detecting transient oscillatory events

We employed widely used spindle and SO detection algorithms (Helfrich et al., 2018; Molle et al., 2011; Staresina et al., 2015) and adjusted them according to the 1/f corrected SO and spindle features for a fully individualized event detection (Hahn et al., 2020).

We detected spindle events (**Figure 3B** & **Figure 3 – figure supplement 1E**) by band-pass filtering the continuous signal ± 2 Hz around the individual spindle peak per electrode. After filtering, we computed the instantaneous amplitude via a Hilbert transform. Next, we smoothed the signal with a running average in a 200 ms window. A sleep spindle was detected, when the signal exceeded the 75-percentile amplitude criterion for a time span of 0.5 to 3 s. We segmented the raw data ± 2.5 s centered on the positive spindle peak.

We detected SO events (**Figure 3 – figure supplement 1F**) by first high-pass filtering the continuous EEG signal at 0.16 Hz and then low-pass filtering at 2 Hz. Based on the filtered signal, we detected the zero-crossings that fulfilled the time criterion (length 0.8 – 2 s). The signal between two consecutive zero-crossings was considered a valid SO if its amplitude exceeded the 75-percentile threshold. We then segmented the raw data ± 2.5 s centered on the negative peak.

### Alleviating power differences

Power differences in the signal can systematically impact cross-frequency coupling measures by changing the signal-to-noise ratio, which in turn influences the precision of the phase estimation of the signal (Aru et al., 2015; Scheffer-Teixeira & Tort, 2016). Because power decreases are apparent across the lifespan (Campbell & Feinberg, 2009, 2016; Hahn et al., 2020; Helfrich et al., 2018), we z-normalized all detected SO and spindle events in the time domain to alleviate this possible confound before calculating phase-amplitude coupling measures (**Figure 3B**).

### Ensuring co-occurrence of SO and sleep spindles

Cross-frequency coupling renders meaningful information of network communication only when the suspected interacting oscillations are present in the signal. Therefore, we only analyzed SO and sleep spindle epochs during which they co-occurred in a 2.5s time window (± ∼2 SO cycles around the spindle peak). Furthermore, we restricted all our coupling analyses to sleep stage NREM3 because of general lower co-occurrence of SO and spindles in NREM2 (**Figure 3 – figure supplement 1CD**), which can cause spurious coupling estimates (Hahn et al., 2020).

### Event-locked cross-frequency coupling

To parameterize the timed coordination between sleep spindles and SO (**Figure 3C**), we computed event-locked cross-frequency coupling analyses (Dvorak & Fenton, 2014; Hahn et al., 2020; Helfrich et al., 2019; Helfrich et al., 2018; Staresina et al., 2015) based on individualized and normalized spindle peak-locked segments. In short, we used a low-pass filter of 2 Hz to extract the underlying SO-component (**Figure 3D**) from the EEG-signal and read out the phase angle corresponding with the sleep spindle peak after applying a Hilbert transform. We then calculated the coupling strength, which is defined as 1 – circular variance using the CircStat Toolbox function circ_r (Berens, 2009) to assess the consistency of the SO sleep spindle interplay.

### Time frequency analyses

We computed event-locked time-frequency representations based on -2 to 2s epochs centered on the negative SO peak (**Figure 3 – figure supplement 1F**). We used a 500 ms Hanning window in 50 ms steps to analyze the frequency power from 5 to 30 Hz in steps of 0.5 Hz. We subsequently baseline corrected the time-frequency representations by z-scoring the data based on the means and standard deviations of a bootstrapped distribution (10000 iterations) for the -2 to -1.5 s time interval of all trials (Flinker et al., 2015; Helfrich et al., 2018).

### Statistical analyses

To compare juggling performance between the *sleep-first* and *wake-first* group and to assess the learning progression, we computed mixed ANOVAS with the between factor condition group (*sleep-first*, *wake-first*) and the repeated measure factor juggling blocks. Because number of juggling blocks differed between adolescents (9, **Figure 2A**) and adults (15, **Figure 2B**) we analyzed the juggling performance separately per age group. Influence of sleep on learning curve (**Figure 2D**) and task proficiency (**Figure 2E**) was assessed by a mixed ANOVA with the between factors condition group (*sleep-first*, *wake-first*) and age group (adolescents, adults) and the repeated factor performance test (pre retention interval 1, post retention interval 1). To correct for multiple comparisons we clustered the data in the frequency (**Figure 3 – figure supplement 1A**), time (**Figure 3B**) and space domain (**Figure 3C & Figure 3 – figure supplement 1B**), using cluster-based random permutation testing (Monte-Carlo method, cluster alpha 0.05, max size criterion, 1000 iterations, critical alpha level 0.05 two-sided; Maris & Oostenveld, 2007). Given our sparse sampling of only 11 scalp electrodes, we set the minimum number of neighborhood electrodes required to be included in the clustering algorithm to zero. For correlational analyses we utilized spearman rank correlations (rho_s_; **Figure 2F** & **Figure 3DE**) to mitigate the impact of possible outliers as well as cluster-corrected spearman rank correlations by transforming the correlation coefficients to t-values (p < 0.05) and clustering in the space domain (**Figure 3DE**). Linear trend lines were calculated using robust regression. To control for possible confounding factors we computed cluster-corrected partial rank correlations (**Figure 3 – figure supplement 3 and 4**). We report partial eta squared (η^2^), Cohen’s d (d) and averaged spearman correlation coefficients (mean rho) as effect sizes. Cluster effect sizes are estimated by first calculating Cohen’s d for every data point in the significant cluster and subsequently averaging across the obtained values.

### Data analyses

We used functions from the Fieldtrip toolbox (Oostenveld et al., 2011), EEGlab toolbox (Delorme & Makeig, 2004), CircStat toolbox (Berens, 2009) and custom written code implemented in MatLab 2015a (Mathworks Inc.) for data analyses. Irregular auto-spectral analysis (IRASA (Wen & Liu, 2016)) was conducted using code obtained from the original research paper.

## DATA AVAILABILITY

The behavioral and electrophysiological preprocessed data and scripts to replicate the main conclusions and figures of the paper are available at https://datadryad.org/stash/share/177ueSz3dyTr3-x6pRaUbZncoOZXNndr_SThSNNkx0A (doi:10.5061/dryad.qfttdz0gh).

## SUPPLEMENTARY FIGURES

**Figure 2 – figure supplement 1.**
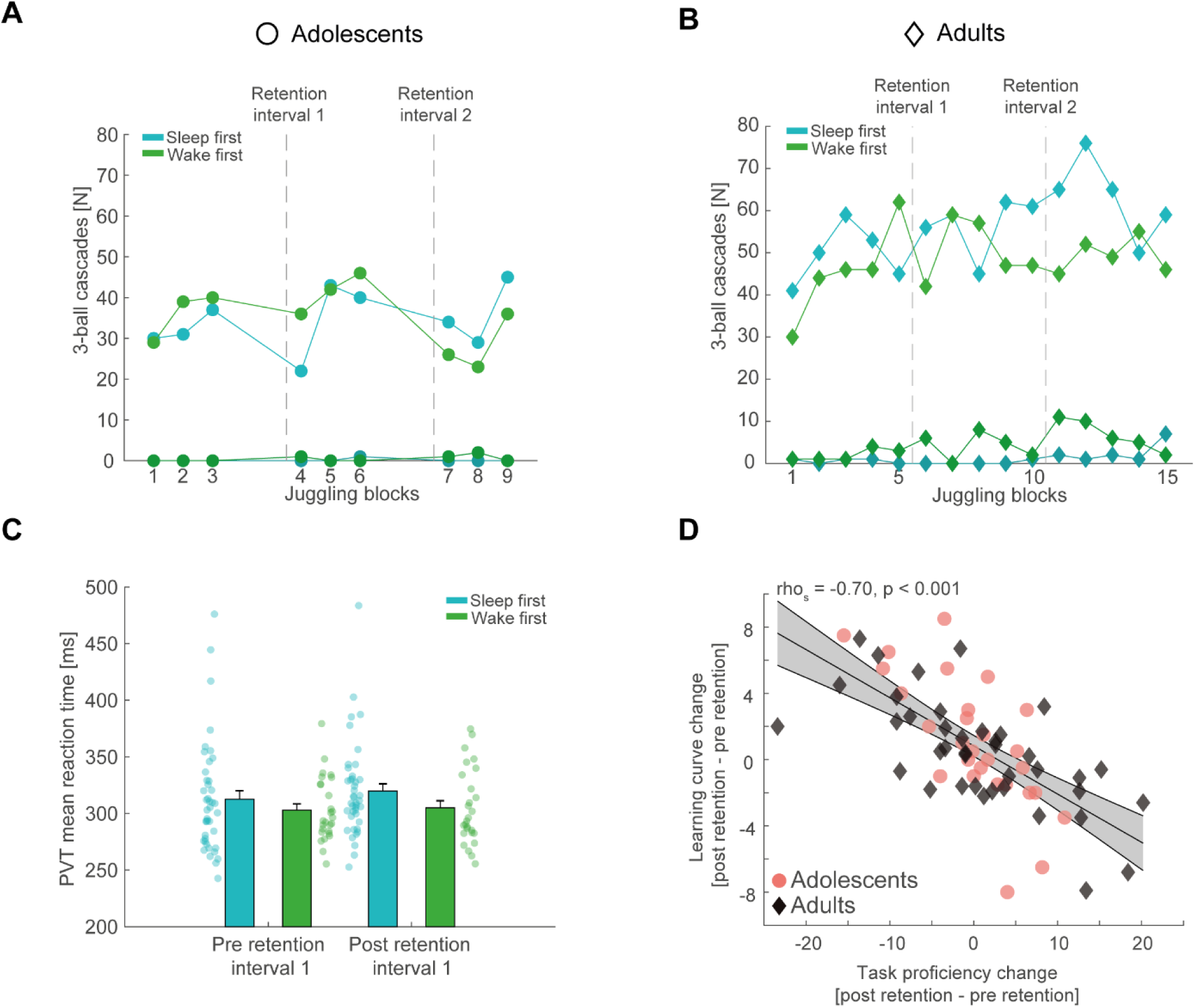
(**A**) Single subject data of successful three-ball cascades per juggling block for well performing adolescents (upper lines) and worse performing adolescents (lower lines) color coded for their respective group affiliation. (**B**) Same conventions as in (A) but for adults. (**C**) Reaction time (mean ± SEM) for the sleep first (blue) and wake first groups (green, collapsed across adolescents and adults) in the psychomotor vigilance tasks conducted before the juggling performance test pre and post the first retention interval. We found no significant difference between the groups (F(1,67) = 1.87, p = 0.18, partial eta² = 0.03) nor between the performance tests (F(1,67) = 1.06, p = 0.31, partial eta² = 0.02). Critically, we found no significant interaction (F(1,67) = 0.35, p = 0.55, partial eta² = 0.01) indicating that participants’ cognitive engagement did not differ in the juggling performance tests due to the preceding sleep or wake intervals. **D**) Spearman rank-correlation between the overnight change in task proficiency (post – pre retention interval) and the overnight change in learning curve with robust linear trend line collapsed over the whole sample after outlier removal. The strong inverse relationship between task proficiency and learning curve originally observed in Figure 2F persisted. Grey-shaded area indicates 95% confidence intervals of the trend line. Adolescents are denoted as red circles and adults as black diamonds.

**Figure 3 – figure supplement 1.**
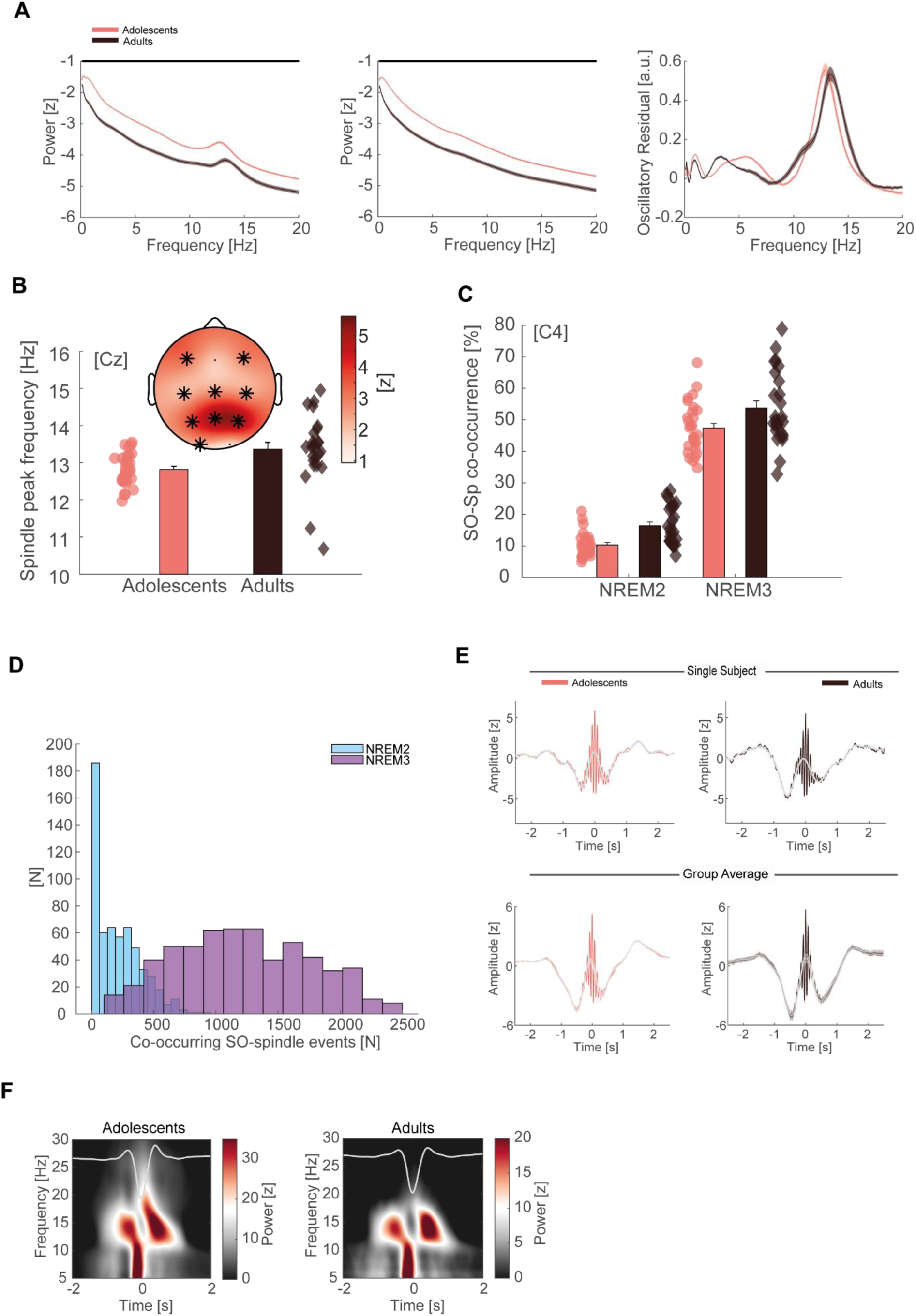
(**A**) Left: Z-normalized EEG power spectra (mean ± SEM) for adolescents (red) and adults (black) during NREM sleep in semi-log space. Data is displayed for the representative electrode Cz unless specified otherwise. Note the overall power difference between adolescents and adults due to a broadband shift on the y-axis. Straight black line denotes cluster-corrected significant differences. Middle: 1/f fractal component that underlies the broadband shift. Right: Oscillatory residual after subtracting the fractal component (A, middle) from the power spectrum (A, left). Both groups show clear delineated peaks in the SO (< 2 Hz) and spindle range (11 – 16 Hz) establishing the presence of the cardinal sleep oscillations in the signal. (**B**) Top: Spindle frequency peak development based on the oscillatory residuals. Spindle frequency is faster at all but occipital electrodes in adults than in adolescents. T-scores are transformed to z-scores. Asterisks denote cluster-corrected two-sided p < 0.05. Bottom: Exemplary depiction of the spindle frequency (mean ± SEM) for adolescents (red) and adults (black) with single subject data points at Cz. (**C**) SO-spindle co-occurrence rate (mean ± SEM) for adolescents (red) and adults (black) during NREM2 and NREM3 sleep. Event co-occurrence is higher in NREM3 (F(1, 51) = 1209.09, p < 0.001, partial eta² = 0.96) as well as in adults (F(1, 51) = 11.35, p = 0.001, partial eta² = 0.18). (**D**) Histogram of co-occurring SO-spindle events in NREM2 (blue) and NREM3 (purple) collapsed across all subjects and electrodes. Note the low co-occurring event count in NREM2 sleep. (**E**) Single subject (top) and group averages (bottom, mean ± SEM) for adolescents (red) and adults (black) of individually detected, for SO co-occurrence-corrected sleep spindles in NREM3. Spindles were detected based on the information of the oscillatory residual. Note the underlying SO-component (grey) in the spindle detection for single subject data and group averages indicating a spindle amplitude modulation depending on SO-phase. (**F**) Grand average time frequency plots (-2 to -1.5s baseline-corrected) of SO-trough-locked segments (corrected for spindle co-occurrence) in NREM3 for adolescents (left) and adults (right). Schematic SO is plotted superimposed in grey. Note the alternating power pattern in the spindle frequency range, showing that SO-phase modulates spindle activity in both age groups.

**Figure 3 – figure supplement 2.**
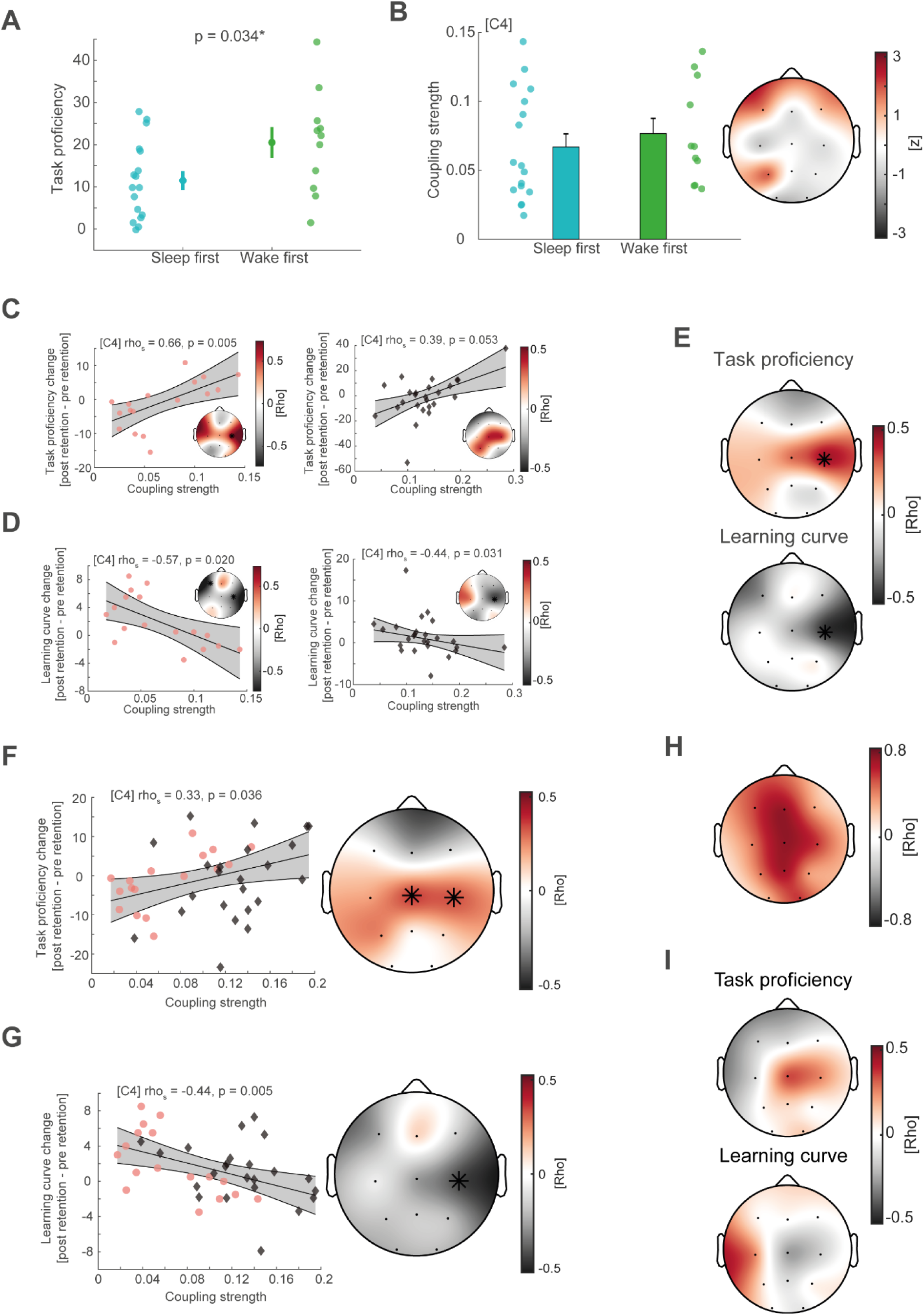
(**A**) Comparison of task proficiency between sleep first and wake first group after the sleep retention interval (mean ± SEM). Adolescents in the wake first group had higher task proficiency given the additional juggling performance test, which also reflects additional training (t(23) = -2.24, p = 0.034). (**B**) Comparison of SO-spindle coupling strength in the adolescent sleep first (blue) and wake first (green) group using cluster-based random permutation testing (Monte-Carlo method, cluster alpha 0.05, max size criterion, 1000 iterations, critical alpha level 0.05, two-sided). Left: exemplary depiction of coupling strength at electrode C4 (mean ± SEM). Right: z-transformed t-values plotted for all electrodes obtained from the cluster test. No significant clusters emerged. (**C**) Left: cluster-corrected correlations between individual coupling strength and overnight task proficiency change (post – pre retention) for adolescents of the sleep-first group with spearman correlation at C4, uncorrected. Asterisks indicate cluster-corrected two-sided p < 0.05. Grey-shaded area indicates 95% confidence intervals of the robust trend line. Participants with a more precise SO-spindle coordination show improved task proficiency after sleep. Right: cluster-corrected correlation of coupling strength and overnight task proficiency change for adults. Independently, adolescents and adults with higher coupling strength have better task proficiency after sleep. (**D**) Left: cluster-corrected correlation of coupling strength and overnight learning curve change for adolescents. Same conventions as in (C). Higher coupling strength related to a flatter learning curve after sleep. Right: Cluster-corrected correlation of coupling strength and overnight learning curve change for adults. Higher coupling strength related to a flatter learning curve after sleep in both age groups. (**E**) Cluster-corrected correlations for coupling strength of co-occurrence corrected events in NREM2 *and* NREM3 sleep with overnight task proficiency change (top) and overnight learning curve change (bottom). Asterisks indicate cluster-corrected two-sided p < 0.05. Similar to our original analyses (Figure 3DE) we found significant cluster-corrected correlations at C4. (**F**) Cluster-corrected correlations between individual coupling strength and overnight task proficiency change (post – pre retention) after outlier removal with spearman correlation at C4, uncorrected. Similar to our original analyses we found a significant central cluster (mean rho = 0.35, p = 0.029, cluster-corrected) after outlier removal. (**G**) Same conventions as in (F) but for overnight learning curve change. Similar to our original analyses we found a significant correlation at C4 (rho = -0.44, p = 0.047, cluster-corrected). (**H**) Topographical plot of spearman rank correlations of coupling strength in the adaptation night and learning night across all subjects. Overall coupling strength was highly correlated between the two measurements (mean rho across all channels = 0.55), supporting the notion that coupling strength remains rather stable within the individual (i.e. trait). (**I**) To investigate a possible state-effect for coupling strength and motor learning, we calculated the difference in coupling strength between the two nights (learning night – adaptation night) and correlated these values with the overnight change in task proficiency and learning curve. We identified no significant correlations with a learning induced coupling strength change. Neither for task proficiency (top) nor learning curve change (bottom).

**Figure 3 – figure supplement 3.**
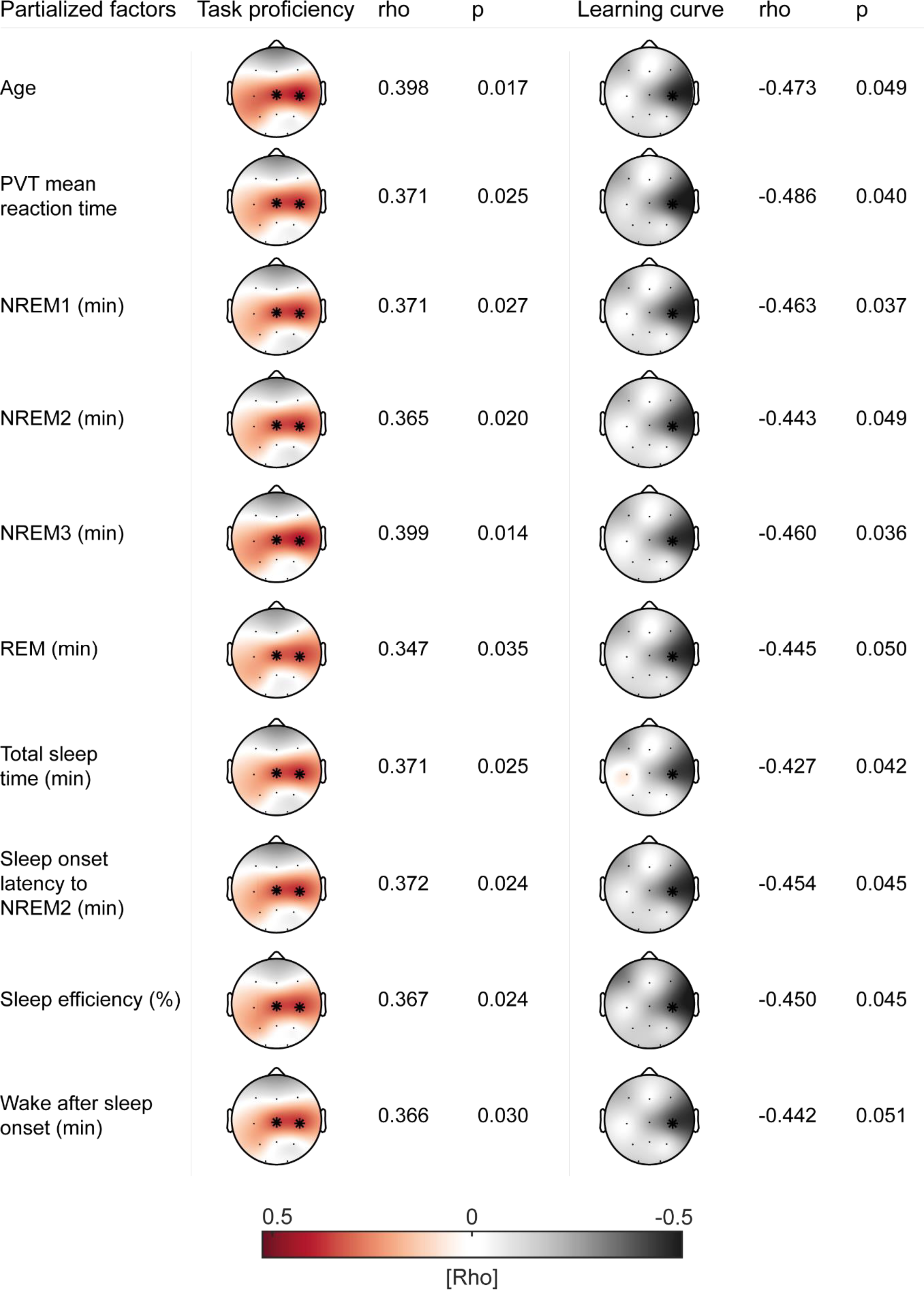
Summary of cluster-corrected partial correlations (Monte-Carlo method, cluster alpha 0.05, max size criterion, 1000 iterations, critical alpha level 0.05, two-sided) of coupling strength with task proficiency (left) and learning curve (right) controlling for possible confounding factors. Asterisks indicate location of the detected cluster. The pattern of initial results remained highly stable.

**Figure 3 – figure supplement 4.**
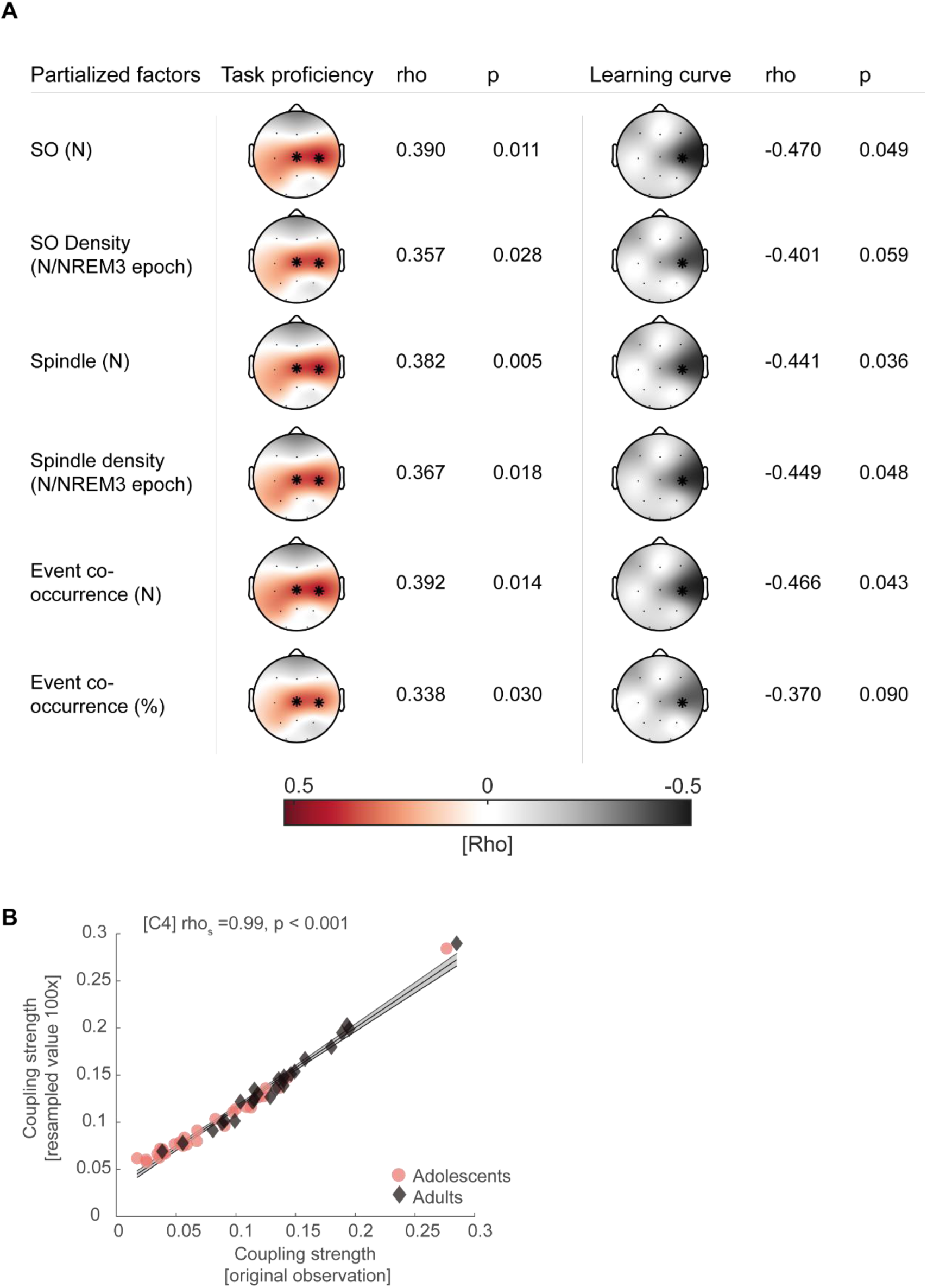
(**A**) Summary of cluster-corrected partial correlations of coupling strength with task proficiency (left) and learning curve (right) controlling SO/spindle descriptive measures at critical electrode C4. Asterisks indicate location of the detected cluster. The pattern of initial results remained highly stable. (**B**) Spearman correlation between resampled coupling strength (N = 200, 100 iterations) and original observation of coupling strength for adolescents (red circles) and adults (black diamonds), indicating that coupling strength is not influenced by spindle event number if at least 200 events are present. Grey-shaded area indicates 95% confidence intervals of the robust trend line.

## SUPPLEMENTARY FILE

**Table 1 related to Figure 1.**
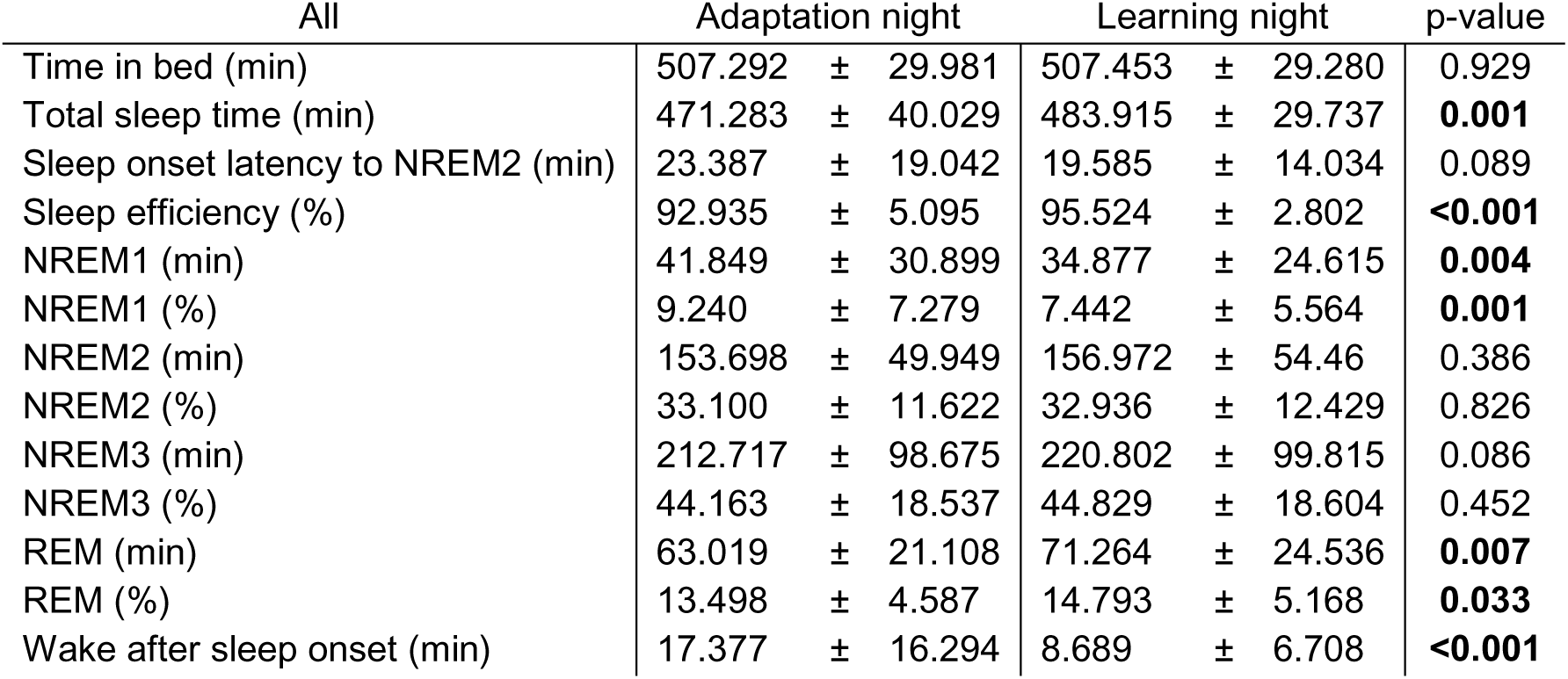
Sleep architecture (mean ± standard deviation) for the adaptation and learning night collapsed across both age groups. Nights were compared using paired t-tests

**Table 2 related to Figure 1.** Summary of sleep architecture and SO/spindle event descriptive measures (at electrode C4) of adolescents and adults across the whole sample (mean ± standard deviation) in the learning night. Independent t-tests were used for comparisons

| All | Adolescents |  | Adults |  | p-value |
| --- | --- | --- | --- | --- | --- |
| Time in bed (min) | 531.679 | ± 18.773 | 480.708 | ± 2.445 | **<0.001** |
| Total sleep time (min) | 506.589 | ± 20.402 | 458.438 | ± 13.550 | **<0.001** |
| Sleep onset latency to NREM2 (min) | 18.821 | ± 15.632 | 20.813 | ± 12.381 | 0.617 |
| Sleep efficiency (%) | 95.520 | ± 2.843 | 95.434 | ± 2.834 | 0.914 |
| NREM1 (min) | 17.839 | ± 8.592 | 55.167 | ± 22.27 | **<0.001** |
| NREM1 (%) | 3.522 | ± 1.686 | 12.100 | ± 5.012 | **<0.001** |
| NREM2 (min) | 124.821 | ± 49.137 | 196.104 | ± 30.804 | **<0.001** |
| NREM2 (%) | 24.728 | ± 9.985 | 42.817 | ± 6.783 | **<0.001** |
| NREM3 (min) | 297.482 | ± 65.336 | 130.792 | ± 43.521 | **<0.001** |
| NREM3 (%) | 58.660 | ± 12.476 | 28.444 | ± 9.230 | **<0.001** |
| REM (min) | 66.446 | ± 27.011 | 76.375 | ± 21.037 | 0.151 |
| REM (%) | 13.090 | ± 5.265 | 16.640 | ± 4.502 | **0.013** |
| Wake after sleep onset (min) | 7.661 | ± 6.285 | 10.125 | ± 7.108 | 0.191 |
| SO (N) | 3499.107 | ± 340.288 | 1855.280 | ± 632.753 | **<0.001** |
| SO density (N/NREM3 epoch) | 6.141 | ± 1.347 | 7.215 | ± 1.997 | **0.025** |
| Spindle number (N) | 3506.571 | ± 742.618 | 1439.080 | ± 580.892 | **<0.001** |
| Spindle density (N/NREM3 epoch) | 5.935 | ± 0.603 | 5.383 | ± 1.159 | **0.032** |
| Event co-occurrence (N) | 1623.357 | ± 295.338 | 753.720 | ± 299.714 | **<0.001** |
| Event co-occurrence (%) | 47.359 | ± 8.030 | 53.745 | ± 11.424 | **0.021** |
| Coupling strength | 0.071 | ± 0.038 | 0.133 | ± 0.0510 | **<0.001** |

**Table 3 related to Figure 2D.** Mixed ANOVA Output comparing juggling learning curves pre- and post retention interval 1 between the condition groups and age groups

| Effect | df | F-statistic | p-value | Effect size ($\eta^2$) |
| --- | --- | --- | --- | --- |
| Performance test (pre-, post retention) | 1 | 1.812 | 0.183 | 0.027 |
| Condition group (sleep first, wake first) | 1 | 0.082 | 0.775 | 0.001 |
| Age group (adolescents, adults) | 1 | 0.992 | 0.323 | 0.015 |
| Condition group*Age group | 1 | 0.238 | 0.627 | 0.004 |
| Performance test*Condition group | 1 | 4.868 | **0.031** | 0.070 |
| Performance test*Age group | 1 | 0.026 | 0.873 | < 0.001 |
| Performance test*Condition group*Age group | 1 | 0.093 | 0.761 | 0.001 |
| Error | 65 |  |  |  |

**Table 4 related to Figure 2E.** Mixed ANOVA Output comparing juggling task proficiency pre- and post retention interval 1 between the condition groups and age groups

| Effect | df | F-statistic | p-value | Effect size ($\eta^2$) |
| --- | --- | --- | --- | --- |
| Performance test (pre-, post retention) | 1 | 0.153 | 0.697 | 0.002 |
| Condition group (sleep first, wake first) | 1 | 0.001 | 0.972 | < 0.001 |
| Age group (adolescents, adults) | 1 | 2.338 | 0.131 | 0.035 |
| Condition group*Age group | 1 | 5.210 | **0.026** | 0.074 |
| Performance test*Condition group | 1 | 1.882 | 0.175 | 0.028 |
| Performance test*Age group | 1 | 0.009 | 0.925 | < 0.001 |
| Performance test*Condition group*Age group | 1 | 0.026 | 0.873 | < 0.001 |
| Error | 65 |  |  |  |

**Table 5 related to Figure 2.** Summary of linear mixed models for predicting learning curve, PVT mean reaction time and task proficiency separately across all performance tests and for the first performance test only using the structure ∼Age group + Time of day + (1|Subjects)

|  | Modeled | Parameter | Beta | T-statistic | df | p-value | Lower-95 CI | Upper-95 CI | Random effects (SD) |
| --- | --- | --- | --- | --- | --- | --- | --- | --- | --- |
| (A) | Learning curve (all performance tests) | Intercept | 2.922 | 6.832 | 202 | < 0.001 | 2.079 | 3.766 | 1.169 |
|  |  | Age group adult | -0.991 | -2.043 | 202 | 0.042 | -1.948 | -0.035 |  |
|  |  | Time of day evening | -1.129 | -2.885 | 202 | 0.004 | -1.901 | -0.357 |  |
| (B) | Learning curve (first performance test) | Intercept | 2.553 | 4.544 | 66 | < 0.001 | 1.431 | 3.675 | 1.648 |
|  |  | Age group adult | -0.540 | -0.945 | 66 | 0.348 | -1.681 | 0.601 |  |
|  |  | Time of day evening | -1.294 | -2.251 | 66 | 0.028 | -2.442 | -0.146 |  |
| (C) | Task proficiency (all performance tests) | Intercept | 14.747 | 5.407 | 202 | < 0.001 | 9.369 | 20.124 | 12.788 |
|  |  | Age group adult | 8.082 | 2.371 | 202 | 0.019 | 1.36 | 14.804 |  |
|  |  | Time of day evening | 2.7467 | 1.999 | 202 | 0.047 | 0.037 | 5.456 |  |
| (D) | Task proficiency (first performance test) | Intercept | 12.804 | 3.791 | 66 | < 0.001 | 6.060 | 19.547 | 9.905 |
|  |  | Age group adult | 6.285 | 1.830 | 66 | 0.072 | -0.571 | 13.141 |  |
|  |  | Time of day evening | 3.470 | 1.004 | 66 | 0.318 | -3.428 | 10.369 |  |
| (E) | PVT mean reaction time (all performance tests) | Intercept | 326.29 | 43.481 | 202 | < 0.001 | 311.5 | 341.09 | 32.550 |
|  |  | Age group adult | -23.762 | -2.585 | 202 | 0.010 | -41.886 | -5.639 |  |
|  |  | Time of day evening | -1.107 | -0.238 | 202 | 0.812 | -10.29 | 8.077 |  |
| (F) | PVT mean reaction time (first performance test) | Intercept | 311.37 | 31.469 | 66 | < 0.001 | 291.61 | 331.12 | 29.015 |
|  |  | Age group adult | -14.209 | -1.413 | 66 | 0.163 | -34.294 | 5.876 |  |
|  |  | Time of day evening | 9.622 | 0.951 | 66 | 0.345 | -10.586 | 29.831 |  |

Table note: linear mixed model we computed with age group (adolescents, adults) and time of day (i.e. performance test in the morning or evening) as fixed effects and subjects as random effects with the fitlme.m matlab function using maximum likelihood estimation. We used reference dummy coding, where the coefficient of the first category is set to 0 (i.e. fixed effect of age group is referenced to adolescents whereas the Time of day fixed effect is referenced to performance tests in the morning).

**Table 6 related to Figure 3DE, Figure 3 – figure supplement 3 & 4.** Summary of sleep architecture and SO/spindle event descriptive measures (at electrode C4) of adolescents and adults in the sleep first group (mean ± standard deviation) in the learning night. Independent t-tests were used for comparisons

| Sleep first | Adolescents |  | Adults |  | p-value |
| --- | --- | --- | --- | --- | --- |
| Time in bed (min) | 530.118 | ± 17.407 | 480.708 | ± 2.445 | **<0.001** |
| Total sleep time (min) | 502.059 | ± 19.204 | 458.438 | ± 13.550 | **<0.001** |
| Sleep onset latency to NREM2 (min) | 21.794 | ± 17.474 | 20.813 | ± 12.381 | 0.834 |
| Sleep efficiency (%) | 94.771 | ± 3.128 | 95.434 | ± 2.834 | 0.484 |
| NREM1 (min) | 16.088 | ± 9.805 | 55.167 | ± 22.27 | **<0.001** |
| NREM1 (%) | 3.206 | ± 1.930 | 12.100 | ± 5.012 | **<0.001** |
| NREM2 (min) | 115.647 | ± 54.532 | 196.104 | ± 30.804 | **<0.001** |
| NREM2 (%) | 23.137 | ± 11.053 | 42.817 | ± 6.783 | **<0.001** |
| NREM3 (min) | 296.147 | ± 71.121 | 130.792 | ± 43.521 | **<0.001** |
| NREM3 (%) | 58.942 | ± 13.758 | 28.444 | ± 9.230 | **<0.001** |
| REM (min) | 74.176 | ± 25.138 | 76.375 | ± 21.037 | 0.763 |
| REM (%) | 14.712 | ± 4.891 | 16.640 | ± 4.502 | 0.200 |
| Wake after sleep onset (min) | 8.294 | ± 7.782 | 10.125 | ± 7.108 | 0.439 |
| SO (N) | 3498.118 | ± 318.304 | 1855.280 | ± 632.753 | **<0.001** |
| SO density (N/NREM3 epoch) | 6.226 | ± 1.508 | 7.215 | ± 1.997 | 0.091 |
| Spindle number (N) | 3477.294 | ± 819.27 | 1439.080 | ± 580.892 | **<0.001** |
| Spindle density (N/NREM3 epoch) | 5.915 | ± 0.669 | 5.383 | ± 1.159 | 0.096 |
| Event co-occurrence (N) | 1617.941 | ± 337.016 | 753.720 | ± 299.714 | **<0.001** |
| Event co-occurrence (%) | 47.709 | ± 8.546 | 53.745 | ± 11.424 | 0.071 |
| Coupling strength | 0.067 | ± 0.039 | 0.133 | ± 0.0510 | **<0.001** |

**Table 7 related to Figure 3 – figure supplement 2AB.** Summary of sleep architecture and SO/spindle event descriptive measures (at electrode C4) of adolescents in the sleep first and wake first group (mean ± standard deviation). Independent t-tests were used for comparisons

| Adolescents | Sleep first |  | Wake first |  | p-value |
| --- | --- | --- | --- | --- | --- |
| Time in bed (min) | 530.118 | ± 17.407 | 534.091 | ± 21.359 | 0.594 |
| Total sleep time (min) | 502.059 | ± 19.204 | 513.591 | ± 21.095 | 0.147 |
| Sleep onset latency to NREM2 (min) | 21.794 | ± 17.474 | 14.227 | ± 11.531 | 0.217 |
| Sleep efficiency (%) | 94.771 | ± 3.128 | 96.677 | ± 1.937 | 0.083 |
| NREM1 (min) | 16.088 | ± 9.805 | 20.545 | ± 5.677 | 0.185 |
| NREM1 (%) | 3.206 | ± 1.930 | 4.009 | ± 1.136 | 0.225 |
| NREM2 (min) | 115.647 | ± 54.532 | 139.000 | ± 37.374 | 0.226 |
| NREM2 (%) | 23.137 | ± 11.053 | 27.187 | ± 7.921 | 0.303 |
| NREM3 (min) | 296.147 | ± 71.121 | 299.545 | ± 58.524 | 0.896 |
| NREM3 (%) | 58.942 | ± 13.758 | 58.224 | ± 10.818 | 0.885 |
| REM (min) | 74.176 | ± 25.138 | 54.500 | ± 26.462 | 0.058 |
| REM (%) | 14.712 | ± 4.891 | 10.583 | ± 5.018 | **0.040** |
| Wake after sleep onset (min) | 8.294 | ± 7.782 | 6.682 | ± 2.831 | 0.518 |
| SO (N) | 3498.118 | ± 318.304 | 3500.636 | ± 387.991 | 0.985 |
| SO density (N/NREM3 epoch) | 6.226 | ± 1.508 | 6.010 | ± 1.109 | 0.687 |
| Spindle number (N) | 3477.294 | ± 819.27 | 3551.818 | ± 641.377 | 0.801 |
| Spindle density (N/NREM3 epoch) | 5.915 | ± 0.669 | 5.965 | ± 0.512 | 0.835 |
| Event co-occurrence (N) | 1617.941 | ± 337.016 | 1631.727 | ± 231.629 | 0.907 |
| Event co-occurrence (%) | 47.709 | ± 8.546 | 46.818 | ± 7.531 | 0.780 |
| Coupling strength | 0.067 | ± 0.039 | 0.077 | ± 0.037 | 0.515 |

